# Arabidopsis hydathodes are sites of intense auxin metabolism and nutrient scavenging

**DOI:** 10.1101/2022.12.01.518666

**Authors:** Jean-Marc Routaboul, Caroline Bellenot, Gilles Clément, Sylvie Citerne, Céline Remblière, Magali Charvin, Lars Franke, Serge Chiarenza, Damien Vasselon, Marie-Françoise Jardinaud, Sébastien Carrère, Laurent Nussaume, Patrick Laufs, Nathalie Leonhardt, Lionel Navarro, Martin Schattat, Laurent D. Noël

## Abstract

Hydathodes are small organs located on the leaf margins of all vascular plants. They release excess xylem sap through guttation when stomata are closed or when the humidity level is high. Many promoter analyses have suggested other hydathode functions in metabolite transport and auxin metabolism, but experimental demonstration is still lacking. Here, we compared the transcriptomic and metabolomic features of mature Arabidopsis hydathodes to the leaf blade. 1460 differentially-expressed genes were identified revealing that genes related to auxin metabolism, transport, stress, DNA, plant cell wall, RNA or wax were on average more expressed in hydathodes. On the other hand, genes involved in glucosinolate metabolism, sulfation pathway, metal handling or photosynthesis were downregulated in hydathodes. In hydathodes, there are an increased expression of auxin transcriptional regulators and biosynthetic genes, a lower expression of auxin transport genes and a differential expression of genes related to its vacuolar storage that is consistent with increased contents of free and conjugated auxin. We also found that ca. 78% of the total content of 52 xylem sap metabolites were removed from guttation fluid at the hydathode level. Using reverse genetics, we showed that the capture of nitrate and phosphate in the guttation fluid relies on the *NRT2.1* and *PHT1;4* transporters, respectively. Thus, hydathodes absorb a significant part of xylem sap nutrients, limiting the loss of valuable chemicals during guttation. Our transcriptomic and metabolomic analyses reveal an organ with its own transcriptomic and physiological identity and highlight hydathode biological processes that may impact the whole plant.

**One sentence summary:** Transcriptome and physiological analysis of mature and healthy hydathodes of Arabidopsis demonstrates that those organs are sites of intense auxin metabolism and nutrient scavenging

## Introduction

A sustained flow of xylem sap from the roots to the shoot supplies both water, organic and inorganic solutes to the shoot of vascular plants. Occasionally, roots may deliver more water than leaves can evaporate when stomata are closed or when the humidity level or atmospheric CO_2_ levels are high. This excess of water could potentially result in a harmful flooding of the leaf intercellular spaces. Prevention of flooding relies on guttation, i.e. the release of the excess of xylem sap at the leaf margin thanks to specialized organs named hydathodes (Feild et al., 2005; Cerutti et al., 2019; Bellenot et al., 2022). Hydathodes are found in all vascular plants and their anatomy has been described in numerous species (Perrin, 1972; Cerutti et al., 2017; Cerutti et al., 2019; Jauneau et al., 2020). This multilayered organ is composed of an outer epidermis containing water pores that resemble stomata but are associated with distinct functions. While stomata regulate the exchange of gases (water vapor, CO_2_ and O_2_), hydathode water pores are specialized in the secretion of liquid. Hydathodes also contain an inner parenchyma called epithem irrigated by a highly branched and hypertrophied xylem system. The number and position of hydathodes are specified very early during leaf morphogenesis and are tightly associated with leaf margin development and leaf serration patterns (Maugarny-Cales and Laufs, 2018).

Although known to botanists for several centuries, our current knowledge on hydathode physiology is very limited. Hydathodes are mostly proposed to be important organs for auxin synthesis and signaling since numerous genes relevant for auxin biosynthesis, transport or signaling are expressed in hydathodes including the auxin reporter *DIRECT REPEAT5* (*DR5*), a synthetic promoter that acts as a readout for auxin activity (Sabatini et al., 1999; Aloni et al., 2003; Alvarez et al., 2009; Eklund et al., 2010; Wang et al., 2011; Muller-Moule et al., 2016). Additionally, auxin itself has also been immuno-localized in Arabidopsis hydathodes (Aloni et al., 2003). The *YUCCA 4* gene important for auxin synthesis and four genes relevant for auxin signaling are more expressed in Arabidopsis hydathodes compared to other leaf tissues (Kim et al., 2021; Yagi et al., 2021). The analysis of multiple mutants of the redundant *YUCCA* biosynthetic genes have shown the importance of auxin synthesis for leaf, veins and hydathode development (Cheng et al., 2006, 2007; Wang et al., 2011; Chen et al., 2014). Many genes coding for transporters are also preferentially expressed in the hydathodes of Arabidopsis, rice or barley (Mudge et al., 2002; Shibagaki et al., 2002; Nazoa et al., 2003; Pilot et al., 2003; Pilot et al., 2004; Nagai et al., 2013; Bailey and Leegood, 2016). Amino acids (Pilot et al., 2004; Bailey and Leegood, 2016) or ions such as NO_3_^-^, PO_4_^-^, or K^+^ (Nagai et al., 2013) are generally less concentrated in the guttation fluid than in the xylem sap suggesting that hydathodes may capture xylem metabolites that have not been used by the leaf (Perrin, 1972; Pilot et al., 2004; Nagai et al., 2013; Bailey and Leegood, 2016). Absorption of ions by the hydathodes would prevent excessive loss of these valuable nutrients during guttation. Some other examples of promoter:*GUS* fusions with hydathode-specific expression profiles are given in Supplemental Table S1.

Healthy hydathodes have been reported to host a rich microbial community (Perrin et al., 1972). Several vascular bacterial pathogens of the *Xanthomonas* and *Clavibacter* genera have evolved to colonize this niche and can infect hydathodes (Lewis and Goodman, 1966; Carlton et al., 1998; Hugouvieux et al., 1998; Cerutti et al., 2017; Bernal et al., 2021; Mullens and Jamann, 2021). In contrast to stomata, hydathode pores seem unable to fully close and may thus facilitate pathogen entry and infection (Cerutti et al., 2017). Yet, immune responses of hydathodes are poorly characterized.

Despite the biological importance of hydathodes, only few molecular, developmental and physiological analyses have been reported. A transcriptomic analysis of macrodissected Arabidopsis hydathodes identified 68 genes which are differentially expressed compared to the leaf blade of three-week-old *in vitro*-grown plants (Yagi et al., 2021). A single cell transcriptomic profiling of Arabidopsis leaf tissues has also found a cluster of 98 genes assigned to the hydathodes (Kim, 2021 #8350}. Thus, the transcriptomic signature of the hydathode tissue remains mostly unexplored.

In this study, a deep transcriptomic description of mature Arabidopsis hydathodes and a detailed metabolomic analysis of guttation fluid were conducted. We revealed that auxin metabolites accumulate at hydathodes in agreement with the increased expression of most genes relevant for the auxin synthesis, transport, storage and signaling. We also demonstrate that two key macronutrient transporter genes strongly expressed in hydathodes limits the loss of nitrate and phosphate in guttation fluid. This study provides a foundation for further physiological studies of hydathodes.

## Results

### Hydathodes possess a transcriptomic signature distinct from the leaf blade

To shed light on the physiological properties of hydathodes, we performed a transcriptomic analysis by RNA sequencing (RNA-Seq) on macrodissected hydathode-enriched samples *versus* leaf blade tissues (comprising mesophyll, stomata, epidermis and some minor veins) of nine-to ten-week-old Arabidopsis plants of Col-0 accession (Figure 1A). Four biological replicates of leaf teeth and leaf blade samples were manually collected using a razor blade. RNA sequencing information about the libraries, number of reads and percentage of raw reads mapped uniquely to the annotated sequences of *Arabidopsis thaliana* chromosomes are given (Supplemental Table S2). The clustering analysis using Euclidean distance showed that hydathode and leaf blade samples clustered in separate groups, indicating distinct transcriptomic signatures (Supplemental Figure S1A). The differential gene expression analysis identified 1460 differentially expressed genes in hydathodes (DEG, FDR<0.001, log_2_ fold change| >0.6), including 610 up-regulated and 850 down-regulated genes (Figure 1B and Supplemental Table S3).

**Figure 1:**
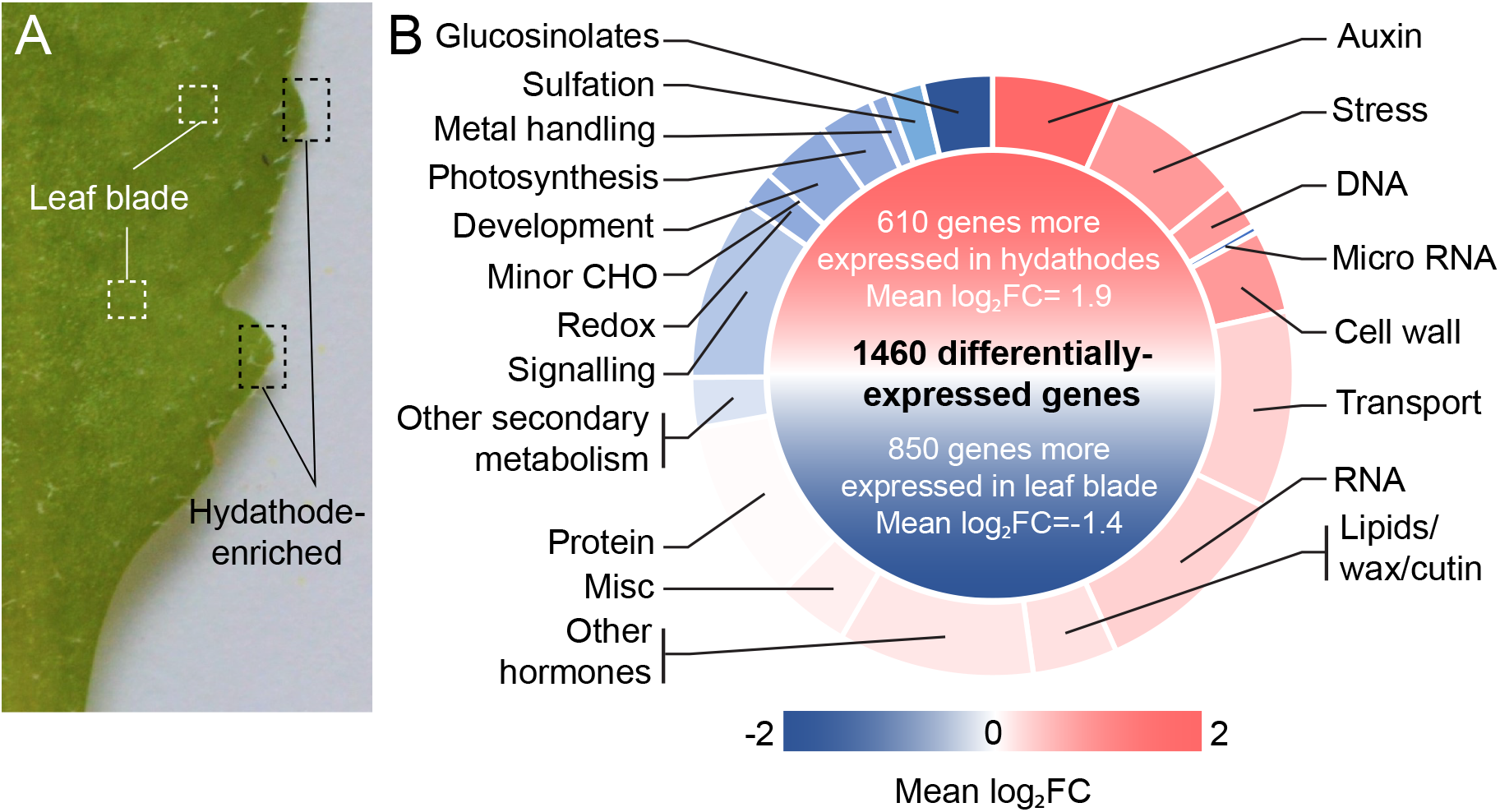
Transcriptomic analysis of hydathode-enriched versus leaf blade samples identifies 1460 differentially-expressed genes (DEG, FDR<0.001) in mature Arabidopsis leaves. (A) Leaf teeth (hydathode-enriched) and leaf blade samples were macro-dissected from nine- to ten-week-old plant leaves and subjected to RNA sequencing. (B) Functional categorization of the 1087 genes differentially-expressed genes with a functional annotation. DEG were mainly annotated using the MapMan analysis classification available from the BAR site (https://bar.utoronto.ca/ntools/cgi-bin/ntools_classification_superviewer.cgi) and expert annotation extracted from literature (Supplemental Table S3). A color key is proportional to the mean log_2_ Fold Change (log_2_FC) of the genes belonging to the given functional category that were globally either more expressed in hydathodes (in red) or in the leaf blade (in blue).

These DEGs were classified by Mapman analysis into functional categories in order to search for enriched functional categories in each sample (Supplemental Figure S1). DEG annotation was manually refined based on the current literature (Supplemental Table S3). DEGs involved in auxin metabolism, stress, DNA, plant cell wall, transport, RNA or lipids were on average more expressed in hydathodes (Figure 1B). On the other hand, glucosinolate synthesis and transport, the sulfation pathway, metal handling or photosynthesis were on average upregulated in the leaf blade. Both auxin- and glucosinolate-related DEG Mapman sub-categories were extracted from the main categories. Consistent with these functional categories, the twenty most DEGs are related to auxin metabolism (i.e. *STY1, STY2, YUC4, YUC5*), transport (*NRT2.1, NIP3;1*), stresses (defensin, disease resistance, chitinase, ABA-induced, CYP714A1) or cell wall (PGL1-pectin lyase-like family, Pollen Ole e 1 like) that are likely marker genes of hydathodes (Table 1). The hydathode transcriptomic signature thus appears distinct from the leaf blade and identifies contrasted biological functions between the hydathode-enriched sample and the leaf blade.

**Table 1:**
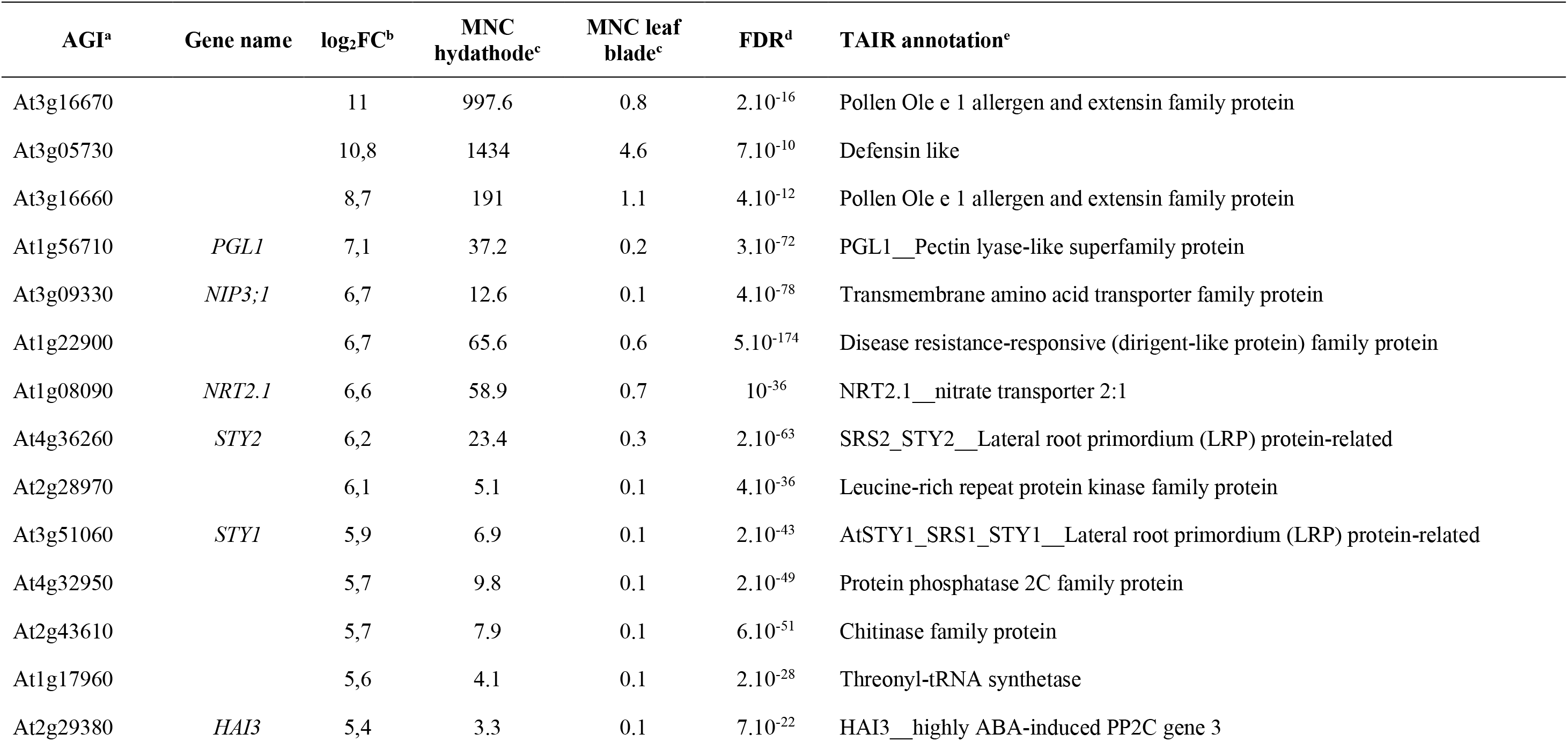
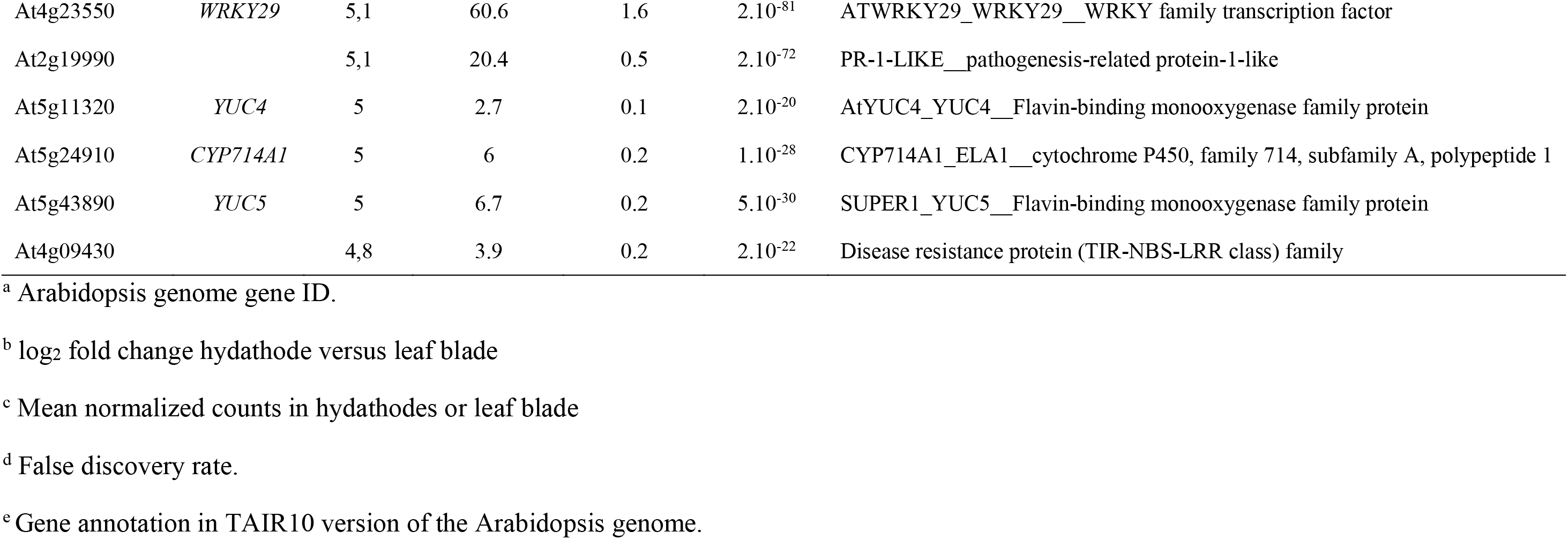
List of the 20 most differentially expressed genes (DEG) in mature Arabidopsis hydathodes relative to the leaf blade (Extracted from Supplemental Table S3).

| AGI<sup>a</sup> | Gene name | log<sub>2</sub>FC<sup>b</sup> | MNC<br>hydathode<sup>c</sup> | MNC leaf<br>blade<sup>c</sup> | FDR<sup>d</sup> | TAIR annotation<sup>e</sup> |
| --- | --- | --- | --- | --- | --- | --- |
| At3g16670 |  | 11 | 997.6 | 0.8 | 2.10<sup>-16</sup> | Pollen Ole e 1 allergen and extensin family protein |
| At3g05730 |  | 10,8 | 1434 | 4.6 | 7.10<sup>-10</sup> | Defensin like |
| At3g16660 |  | 8,7 | 191 | 1.1 | 4.10<sup>-12</sup> | Pollen Ole e 1 allergen and extensin family protein |
| At1g56710 | *PGL1* | 7,1 | 37.2 | 0.2 | 3.10<sup>-72</sup> | PGL1__Pectin lyase-like superfamily protein |
| At3g09330 | *NIP3;1* | 6,7 | 12.6 | 0.1 | 4.10<sup>-78</sup> | Transmembrane amino acid transporter family protein |
| At1g22900 |  | 6,7 | 65.6 | 0.6 | 5.10<sup>-174</sup> | Disease resistance-responsive (dirigent-like protein) family protein |
| At1g08090 | *NRT2.1* | 6,6 | 58.9 | 0.7 | 10<sup>-36</sup> | NRT2.1__nitrate transporter 2:1 |
| At4g36260 | *STY2* | 6,2 | 23.4 | 0.3 | 2.10<sup>-63</sup> | SRS2_STY2__Lateral root primordium (LRP) protein-related |
| At2g28970 |  | 6,1 | 5.1 | 0.1 | 4.10<sup>-36</sup> | Leucine-rich repeat protein kinase family protein |
| At3g51060 | *STY1* | 5,9 | 6.9 | 0.1 | 2.10<sup>-43</sup> | AtSTY1_SRS1_STY1__Lateral root primordium (LRP) protein-related |
| At4g32950 |  | 5,7 | 9.8 | 0.1 | 2.10<sup>-49</sup> | Protein phosphatase 2C family protein |
| At2g43610 |  | 5,7 | 7.9 | 0.1 | 6.10<sup>-51</sup> | Chitinase family protein |
| At1g17960 |  | 5,6 | 4.1 | 0.1 | 2.10<sup>-28</sup> | Threonyl-tRNA synthetase |
| At2g29380 | *HAI3* | 5,4 | 3.3 | 0.1 | 7.10<sup>-22</sup> | HAI3__highly ABA-induced PP2C gene 3 |
| At4g23550 | *WRKY29* | 5,1 | 60.6 | 1.6 | $2.10^{-81}$ | ATWRKY29_WRKY29__WRKY family transcription factor |
| At2g19990 | | 5,1 | 20.4 | 0.5 | $2.10^{-72}$ | PR-1-LIKE__pathogenesis-related protein-1-like |
| At5g11320 | *YUC4* | 5 | 2.7 | 0.1 | $2.10^{-20}$ | AtYUC4_YUC4__Flavin-binding monooxygenase family protein |
| At5g24910 | *CYP714A1* | 5 | 6 | 0.2 | $1.10^{-28}$ | CYP714A1_ELA1__cytochrome P450, family 714, subfamily A, polypeptide 1 |
| At5g43890 | *YUC5* | 5 | 6.7 | 0.2 | $5.10^{-30}$ | SUPER1_YUC5__Flavin-binding monooxygenase family protein |
| At4g09430 | | 4,8 | 3.9 | 0.2 | $2.10^{-22}$ | Disease resistance protein (TIR-NBS-LRR class) family |

<sup>a</sup> Arabidopsis genome gene ID.
<sup>b</sup> log<sub>2</sub> fold change hydathode versus leaf blade
<sup>c</sup> Mean normalized counts in hydathodes or leaf blade
<sup>d</sup> False discovery rate.
<sup>e</sup> Gene annotation in TAIR10 version of the Arabidopsis genome.

### Independent validations of the hydathode sampling and transcriptomic results

In order to verify the robustness of our transcriptomic results, we first compared published reports of promoter:*GUS* reporter lines with our results (Supplemental Table S1). Out of 41 promoter:*GUS* reporter lines which display preferential GUS activity in hydathodes, 33 showed significant and positive log_2_FC in our transcriptomic analysis (Average log_2_FC=2.0). Reciprocally, out of 12 promoter:*GUS* reporter lines which display preferential GUS activity in the leaf blade, 11 showed significant and negative log_2_FC in our transcriptomic analysis (Average log_2_FC=-1,4). These genes are for instance relevant for glucosinolate metabolism (Schuster et al., 2006; Gigolashvili et al., 2007; Redovnikovic et al., 2012) or glutamine transport (Pratelli et al., 2010). Because most of these studies were conducted on young *in vitro*-grown seedlings, we studied the expression of several DEG thanks to existing promoter:*GUS* reporter lines in eight-week-old plants. In hydathodes, the *YUCCA 2, 5, 8* and *9* genes as well as the *TAA1* auxin biosynthesis gene and *PHT1;4* phosphate transporter gene were all intensely expressed in mature hydathodes (Supplemental Figure S2). These results are consistent with our transcriptomic results obtained at the same plant age. In addition, the *DR5:GUS* reporter was specifically active at the tip of the hydathodes compared to *YUCCA* and *TAA1* genes (Supplemental Figure S2), consistent with previous reports (Aloni et al., 2003) and higher auxin signaling at hydathodes (Supplemental Table S3). Observation of the *DR5:VENUS* reporter showed that strong auxin signaling was not uniform throughout the hydathode epithem but was higher in the epithem region close to the vasculature (Supplemental Figure S3). In contrast to these genes preferentially expressed in hydathodes, *GDU4.7* encoding the glutamine vascular transporter (Pratelli et al., 2010) is expressed in the vasculature but not in the hydathodes in agreement with our transcriptomic results (Supplemental Table S3 and Supplemental Figure S2).

We also compared our results with previous transcriptomic studies, which identified 68 Arabidopsis genes more expressed in hydathodes relative to the leaf tissues surrounding the hydathode and a single cell transcriptomic cluster of 98 genes putatively corresponding to hydathode cell types (Kim et al., 2021; Yagi et al., 2021). Sixty five out of 68 and 47 out of 98 were also differentially enriched in our hydathode samples, respectively. Our hydathode samples also shared 74%, 45% and 39% of DEGs with the clusters corresponding to guard cells, epidermis and vasculature which are components of the hydathode organ (Kim et al., 2021). In contrast, mesophyll parenchyma shared only 15% of DEGs more expressed in the hydathode. Last but not least, novel promoter:*GUS* reporter transgenic lines were constructed using the 2-kb region upstream of the ATG of four genes amongst the most differentially expressed genes: AT3G16670 (Pollen OLE 1), AT3G05730 (defensin-like), AT1G62510 (lipid transfer protein) and AT1G56710 (PGL1: Pectin Lyase-like family) (Table 1, Supplemental Table S3). All four promoters were specifically active in hydathodes (Figure 2). Three were specifically restricted to this organ in leaves while AT1G62510 also exhibited a weak and diffuse expression in leaf blade.

**Figure 2:**
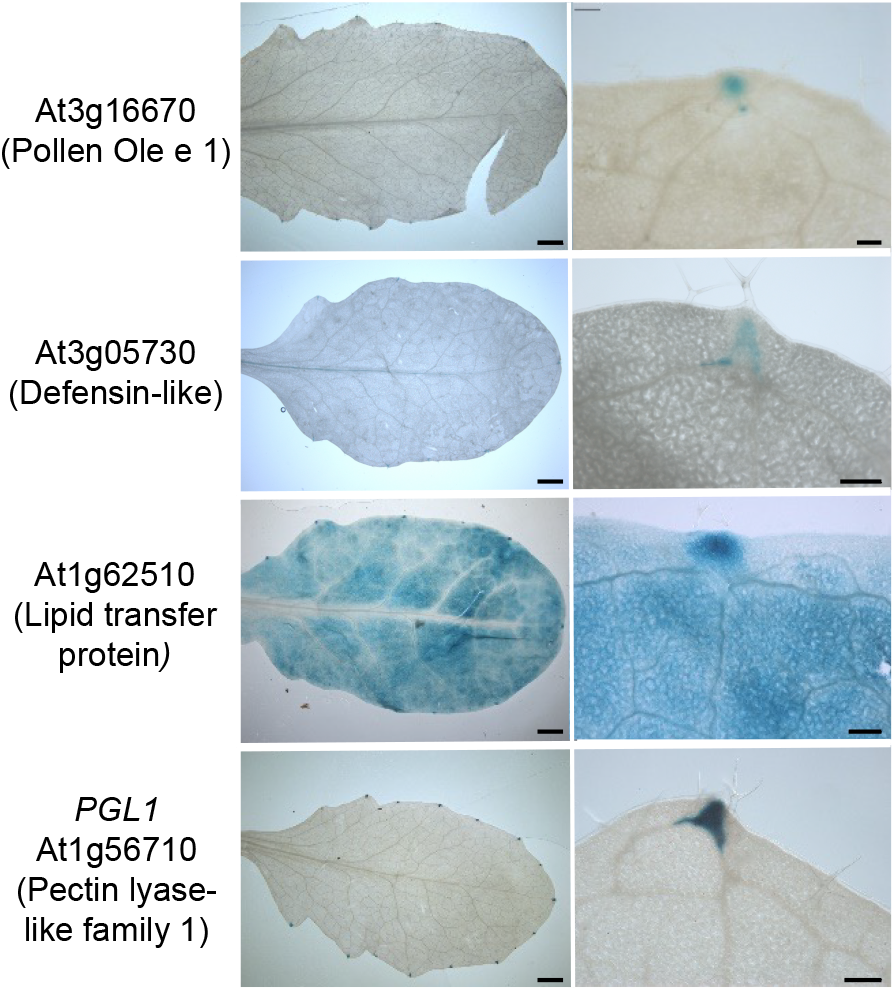
Detection of GUS activity (blue) in leaves of eight-week-old transgenic plants carrying promoter:*GUS* reporter fusions for *Pollen Ole e 1* (*At3g16670*), *Defensin-like* (*At3g05730*), *Lipid transfer protein* (*At1g62510*) and *Pectin lyase-like family PGL1* (*At1g56710*). Scale bars = 2 mm (Left panels), 200 μm (Right panels).

Altogether, these results indicate that the hydathode samples were indeed enriched in hydathode tissue and that our transcriptomic results are valid. We thus mined this dataset to get further insights into hydathode physiology.

### Hydathodes are major sites for auxin metabolism, storage, transport and signaling

Because many genes important for auxin metabolism were reported to be expressed in hydathodes (Supplemental Table S3 for some examples), we measured the accumulation of free auxin by mass spectrometry in hydathodes and leaf blades (Figure 3A). Thirty seven percent more free auxin was measured in hydathodes. Such auxin gradient could be established by differential auxin synthesis and/or transport. The transcriptomic dataset shows that 42/67 genes relevant for auxin metabolism were upregulated in hydathodes (Figure 3B). Those genes encode key biosynthetic enzymes such as TAA1, AO1 or six YUCCA (Average log_2_FC=3.2) and positive regulators from the NGA, SHI or TCP transcription factor families (Average log_2_FC 2.5, Figure 3B and Supplemental Table S3). Therefore, hydathodes are definitely organs of active auxin synthesis in mature leaves. In contrast, auxin transport seems globally repressed. Genes encoding auxin efflux carriers PIN-formed *PIN1* (basipetal transport, Galweiler et al., 1998) or *PIN5* (cytosol to ER transport, Mravec et al., 2009) were less expressed in hydathodes (Figure 3C). Their positive regulators *GOLVEN SECRETORY PEPTIDE GLV1* and *GLV2* and *PINOID-BINDING PROTEIN* (*PBP1*) are also less expressed in hydathodes (Benjamins et al., 2003; Whitford et al., 2012; Ghorbani et al., 2016; Barbosa et al., 2018) while their negative regulator *MACCHI-BOU4* (*MAB4*) is more expressed (Furutani et al., 2014). Parallel to PIN-mediated transport, the *LAX2* transporter gene from the *AUX/LAX* family, known to be expressed in the leaf vasculature except along the leaf margin, was less expressed in hydathodes (Kasprzewska et al., 2015). The ATP-binding cassette (ABC) *ABCB14*, which promotes local influx of auxin was upregulated in hydathodes (Kaneda et al., 2011). The vaculolar storage of auxin at hydathodes could also be promoted by both increasing expression of two IAA-amido synthetase *GH3* that conjugate and inactivate excess IAA to amino acids (Staswick et al., 2005) and by reducing expression of *WALLS ARE THIN1* (*WAT1*) that encodes a tonoplastic transporter important for auxin export (Ranocha et al., 2013). We measured auxin storage capability by quantifying oxindole-3-AIA, which is a down-product of this inactivation pathway (Hayashi et al., 2021). Oxindole-3-AIA was 22% more abundant in hydathodes compared to the leaf blade samples (Figure 3C). Finally, we also observed a specific auxin signaling and response in the hydathode. Two main families of transcription factors regulate the expression of auxin response genes (Guilfoyle and Hagen, 2007). Auxin Response factors (ARFs) are repressed by AUXIN/INDOLE ACETIC ACID (Aux/IAA) proteins. In response to auxin, the degradation of Aux/IAAs by the proteasome releases the ARFs that bind to the auxin-responsive cis-acting element of early auxin response gene promoter, including Aux/IAAs, small auxin-up RNAs (SAURs) and GH3 proteins. We found that IAA28 and IAA29 and ARF 3, 4 and 18 were more expressed in hydathodes (Figure 3A). In addition, two other transcription factors WRKY57 and MYB77 were also differentially expressed. WRKY57 interacts with IAA29 and is regulated by auxin (Jiang et al., 2014). MYB77 interacts with ARF to increase auxin response (Shin et al., 2007). AIR3, SAUR 17, SAUR32 and SAUR36 or Big Grain 1 (BIG1) have also been shown to be activated in response to auxin (Xie et al., 2000, Hou, 2013 #8423; Mishra et al., 2017).

**Figure 3:**
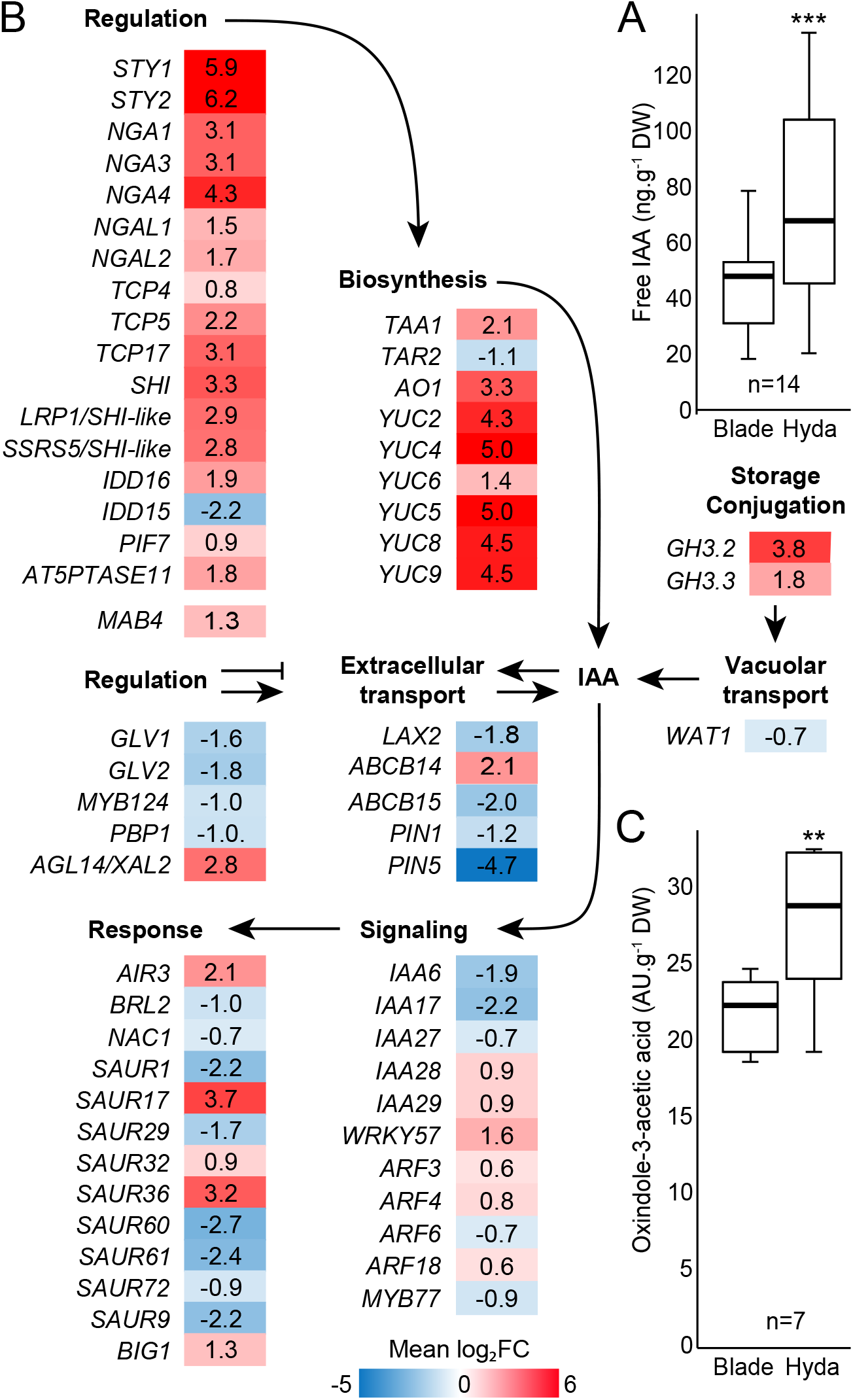
Hydathodes are important organs for auxin metabolism and accumulation in mature Arabidopsis leaves. (A) Boxplot representation of concentrations of free indole-acetic acid (IAA) in leaf blade (Blade) and hydathodes (Hyda) of eight-week-old plants were determined by GC-MS. (B) Expression ratio (Hydathodes vs. leaf blade) of differentially-expressed genes (DEG, Table S3) important for auxin biology, including biosynthetic genes and their regulators, transporter genes and their regulators as well as their downstream signaling components and auxin-responsive genes. Genes more expressed in hydathodes are shown in red, those more expressed in leaf blade in blue. Numbers refer to mean log_2_ fold change (log_2_FC). (C) Boxplot representation of concentrations of auxin conjugate precursor oxindole-3-acetic acid in leaf blade (Blade) and hydathodes (Hyda) of eight-week-old plants were determined by GC-MS. Statistically significant differences were determined using a paired *t*-test (**, p<0.01; ***, p<0.001). n= number of biological replicates.

Altogether, our transcriptomic analysis is consistent with hydathodes being sites of auxin synthesis and signaling as a result of increased auxin synthesis, lower basipetal transport and higher vacuolar storage.

### Transporters expressed in hydathodes prevent excessive loss of metabolites during guttation

According to our transcriptome analysis of hydathode and leaf blade tissues, over 64 transporter genes are preferentially expressed in hydathodes (Table 2). These include genes coding for transporters of water, ions (nitrate, phosphate, sulfate, calcium, zinc, iron, copper, chloride, boron arsenate), hormones (ABA, GA, auxin, cytokinins), sugars, peptides, waxes and other organic compounds. This is in agreement with previous observations showing that genes encoding transporters for nitrate (*NRT1.1* and *NRT2.1*) and inorganic phosphate (Pi, *PHT1;4*) are expressed in hydathodes (Nazoa et al., 2003; Misson et al., 2004; Krouk et al., 2010).

**Table 2:**
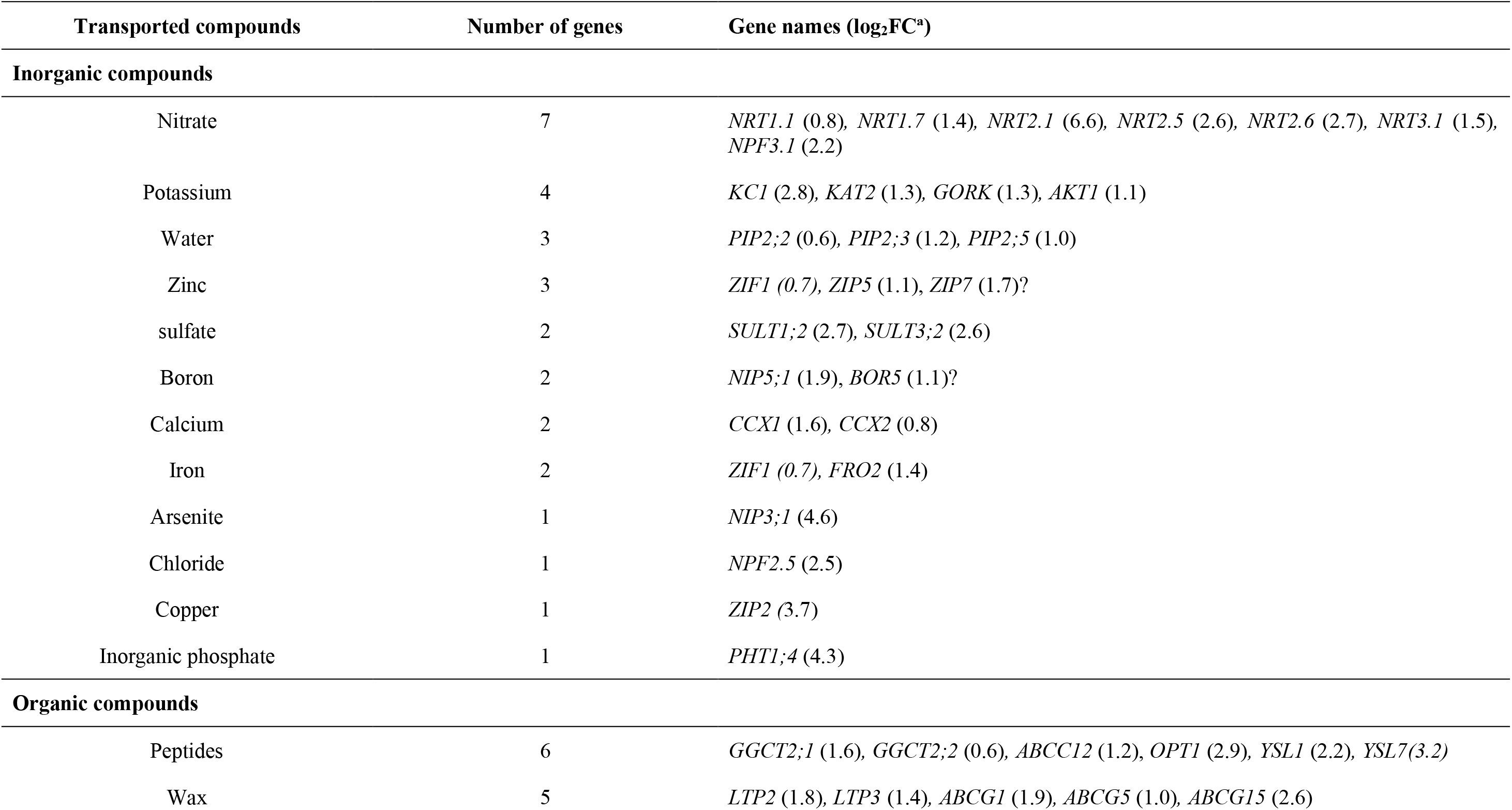
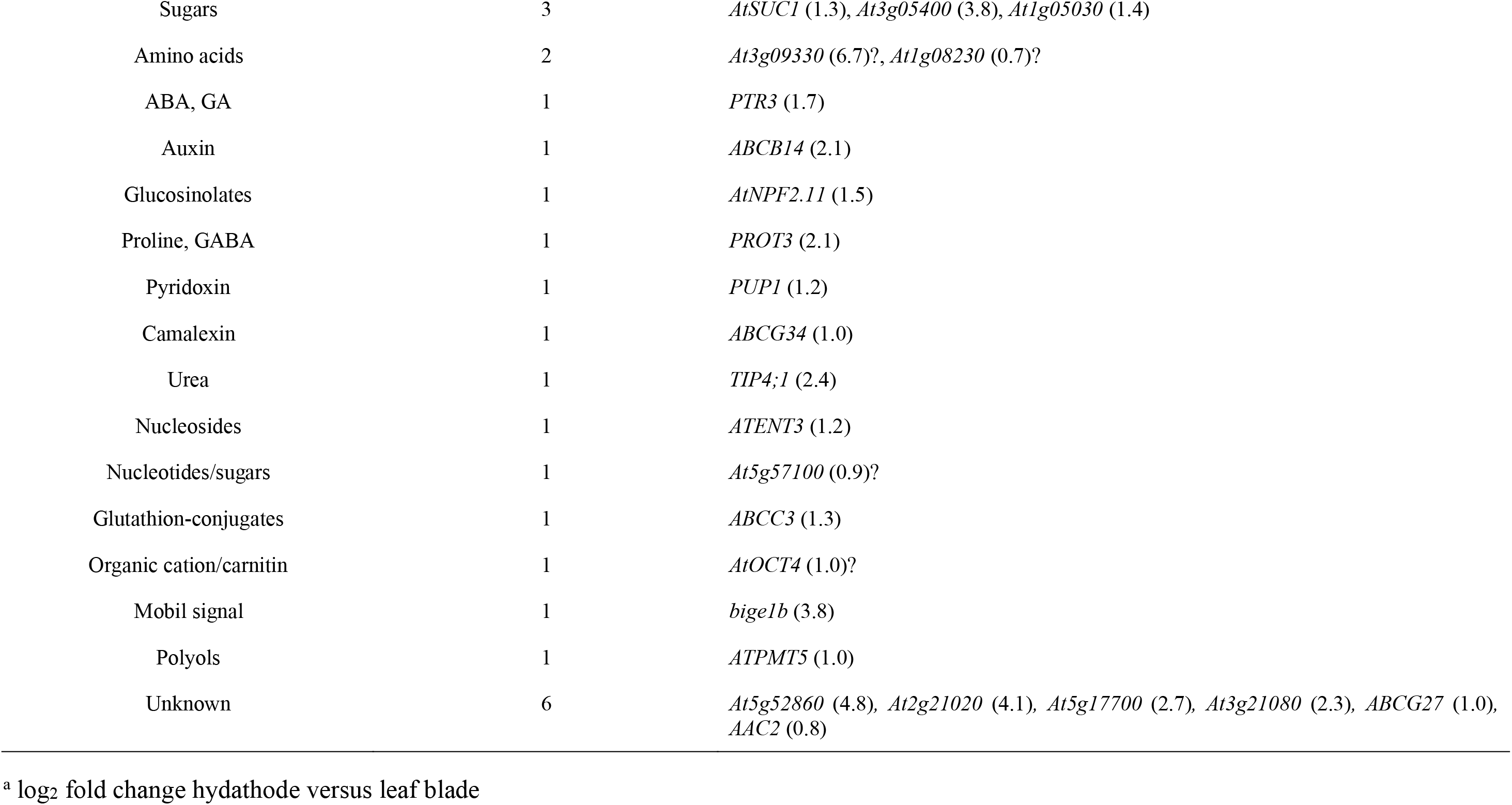
List of the genes encoding transporters which are overexpressed in mature Arabidopsis hydathodes relative to the leaf blade (Extracted from Supplemental Table S3).

These observations suggest that transporters inside hydathodes could actively modify guttation fluid composition. In order to test this hypothesis, guttation was sampled at the border of a half leaf from which the margin was excised, as a proxy of a pre-hydathode xylem sap sample. Its composition was compared by GC-MS to the genuine guttation fluid from the intact other half leaf and xylem sap exuding from the petiole (Figure 4A). Only 9% and 2% of the initial metabolites relative to the petiole sample are remaining in the pre-hydathode and guttation fluid samples, respectively. This indicates that both the leaf tissues and the hydathodes capture metabolites from the xylem sap. The concentrations of 23 metabolites (including 13 amino acids, 5 organic acids, 4 sugars and myo-inositol, p<0.05) in the guttation fluid were significantly lower than in the pre-guttation samples (Figure 4B). A significant uptake of Pi and nitrate in the hydathodes was also measured (Figure 4C and D). Lower concentrations of phosphorus, calcium and magnesium but not ammonium nor potassium were measured by ICP-MS in the guttation fluid relative to the pre-hydathode fluid (Figure 4E). These results indicate that the hydathodes significantly and specifically retains given organic and mineral compounds before guttation occurs.

**Figure 4:**
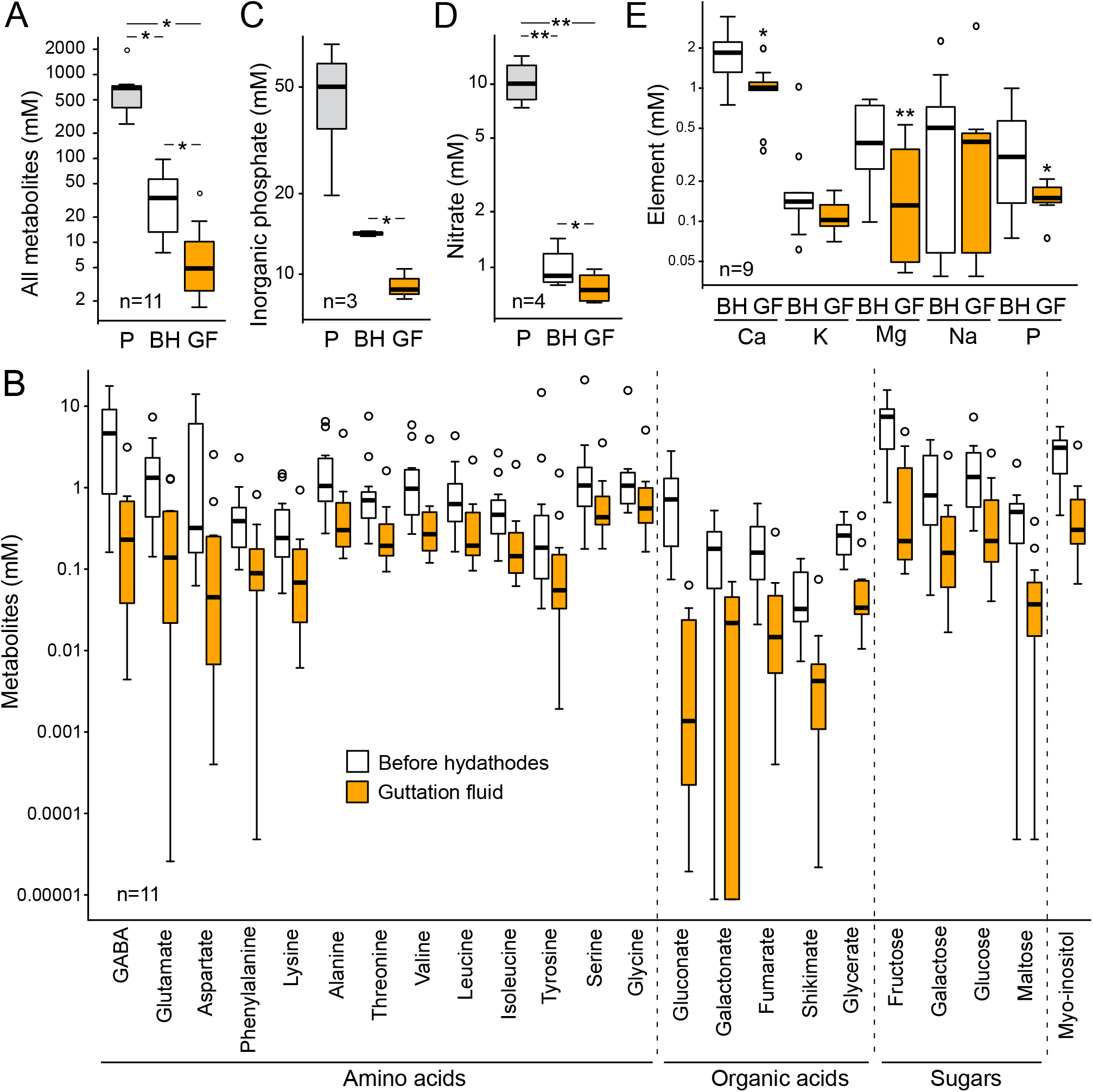
Metabolomic analysis of fluids collected at the petiole (P), before hydathodes (BH) or at the leaf margin (guttation fluid, GF) of leaves from four-week-old plants. (A, B) Concentrations of 52 metabolites were determined by GC-MS in P, BH and G fluids (Table S4). (A) Boxplot representation of total metabolite concentrations in the three types of samples. (B) Box plot representation of 23 out of the 52 metabolites which are in significantly lower concentrations in the guttation fluid (GF) compared to the fluids sampled before hydathodes (BH). Statistically significant differences were determined using non-parametric Kruskal-Wallis test (p<0.05). (C) Boxplot representation of nitrate concentrations in P, BH and G fluids. (D) Boxplot representation of inorganic phosphate concentrations in P, BH and G fluids. (E) Boxplot representation of mineral elements concentrations in P, BH and G fluids measured with ICP-MS. (C,D) Statistically significant differences were determined using a paired *t*-test (*, p<0.05; **, p<0.01). n= number of biological replicates.

In order to determine the biological significance of transporters in guttation fluid composition, the guttation fluid composition of the *nrt2.1* and *pht1;4* mutants was determined for nitrate and Pi, respectively. Higher concentrations of nitrate and Pi were measured in the guttation fluid of *nrt2.1* and *pht1;4* mutants compared to wild-type plants, respectively (Figure 5). This indicates that *NRT2.1* and *PHT1;4* contribute actively to the scavenging of nitrate and Pi from the guttation fluid, respectively. The contribution of the other transporters to the retrieval of the other substrates remains to be determined.

**Figure 5:**
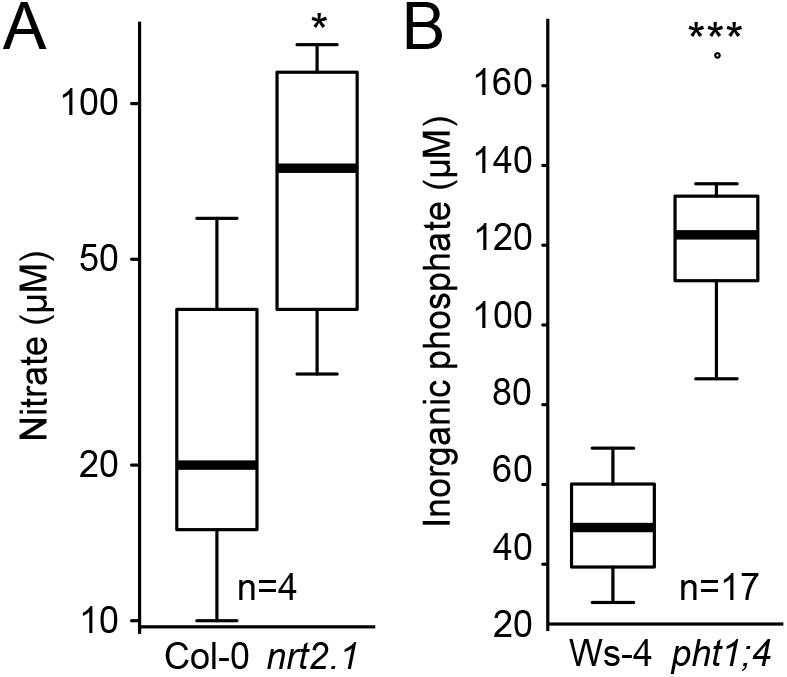
NRT2.1 nitrate and PHT1;4 phosphate transporters are important to capture nitrate and phosphate in hydathodes, respectively. Boxplot representations of nitrate (A) and inorganic phosphate (B) concentrations measured in guttation fluids of four-week-old *nrt2.1* (N859604) and *pht1;4-1* mutants relative to their wild-type controls Col-0 and Ws-4, respectively. Statistically significant differences were determined using a paired *t*-test (*, p<0.05; ***, p<0.001). n= number of biological replicates.

Taken together, those results demonstrate that Arabidopsis hydathodes are active sites of transporter-mediated scavenging of small organic and inorganic compounds, which limits the loss of valuable nutrients during guttation.

### Several physiological processes contributing positively to immunity are dampened in hydathodes

The transcriptomic data also indicates that genes relevant to plant constitutive (cuticular wax and cutin metabolism, cell wall modification and degradation, inhibitory chemicals or proteins synthesis) and induced defenses (glucosinolate synthesis, hormone crosstalks) were differentially expressed in the hydathodes as compared to the leaf blade, as illustrated for instance with the significant higher number of DEGs related to the “Stress” functional category (Supplemental Figure S1, Table 3 and Supplemental Table S3).

**Table 3:**
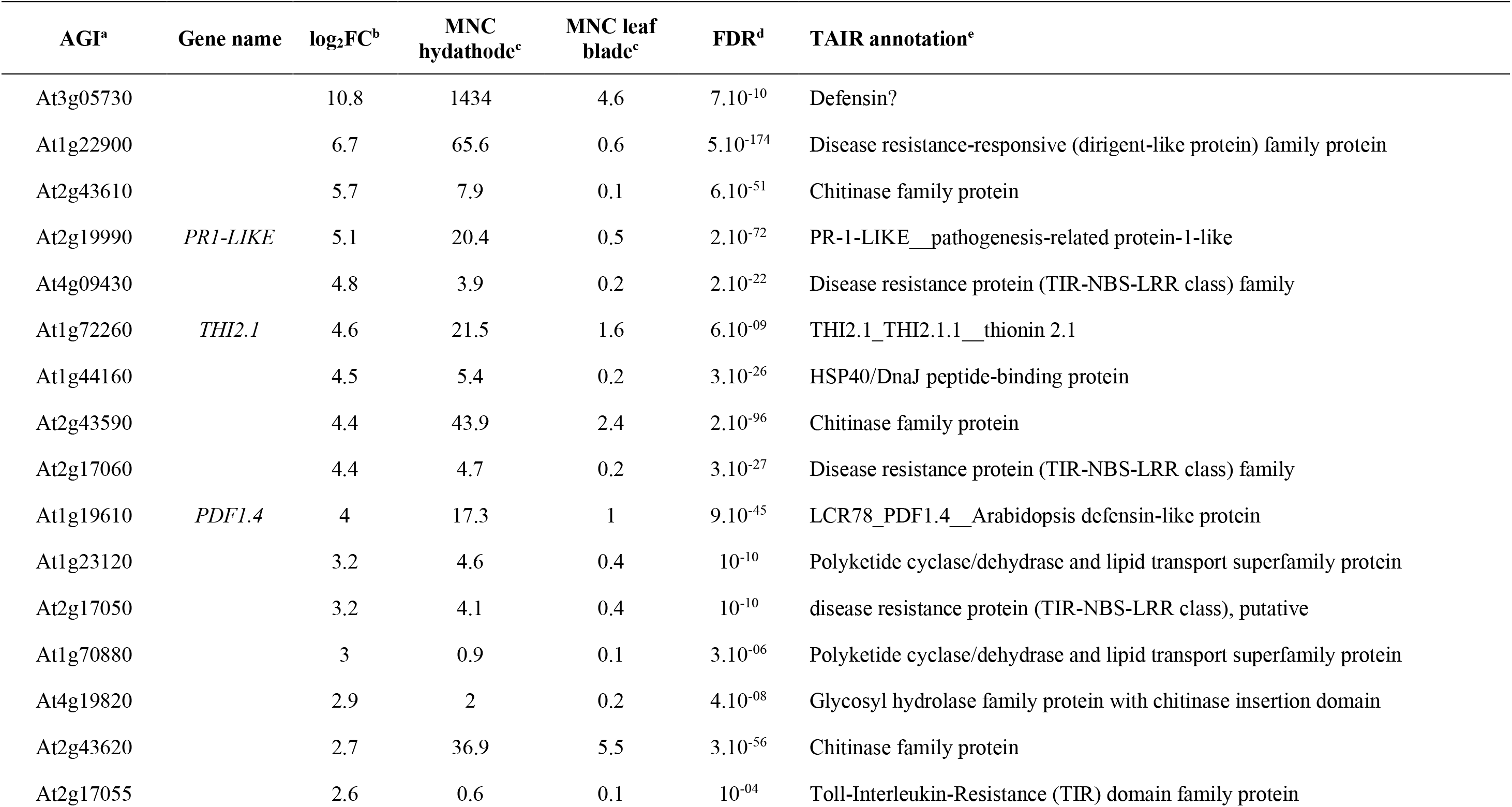
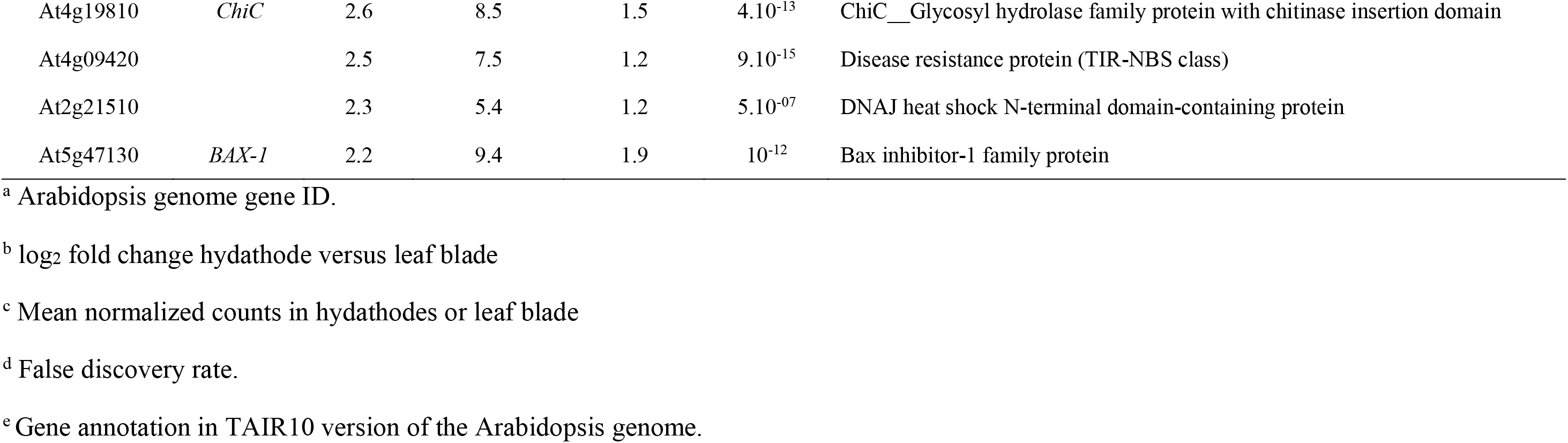
List of the defense-related 20 most differentially expressed genes (DEG) in mature Arabidopsis hydathodes relative to the leaf blade (Extracted from Supplemental Table S3).

#### Wax and cutin metabolism

The functional category “lipid metabolism” was significantly enriched in our dataset (Figure 1B and Supplemental Figure S1C) and a significant number of genes related to lipid signaling, waxes and cutins synthesis and transport were found to be more expressed in hydathodes (Supplemental Table S3). Most glycerolipid-related genes were downregulated, such as *MGDC* that is also a marker of phosphate starvation (Kobayashi et al., 2009). The most differentially expressed gene was *WSD1* with a log_2_FC of 4.2. WSD1 drives the conversion of primary alcohols into esters and is expressed in hydathodes (Li et al., 2008). Waxes and cutins that compose the cuticle are composed of five classes, i.e. acids, aldehydes, alcohols, alkanes, ketones and esters. In contrast to the stem, the leaf only contains minute amounts of esters (Patwari et al., 2019). This increased *WSD1* expression in hydathodes could thus favor the accumulation of wax esters, similar to the one observed in stems. Unfortunately, analysis of the hydathode wax ester would require large amounts of plant material that are for now not adapted to the small hydathode size. Lipophilic cuticles on leaf surfaces represent a protection against water loss or microbes. It would thus be interesting to investigate to what extent these putative wax modifications around the hydathodes might facilitate or impair infection by microbes.

#### Cell wall degradation and modification

We examined the 44 cell wall-related DEGs, refining the annotation with Arabidopsis literature (Supplemental Table S3). Three *Pectin Lyase-like* (*PLs*) and four *Pectin Methyl Esterase inhibitors* (*PMEIs*) genes are upregulated in hydathodes when the *MEPCRA Pectin Methyl Esterase* (*PME*) (Coculo and Lionetti, 2022) and *Pectin Acetyl Esterase* (*PAE2*) are all down regulated. Pectins constitute up to ~50% of Arabidopsis leaf walls (Zablackis et al., 1995) and are especially important in the middle lamella, acting as glue between the walls of two adjacent cells thus preventing the cells from separating or sliding against each other (Zamil and Geitmann, 2017). PME catalyzes the demethoxylation of pectin and thus favors cell to cell adhesion through the formation of calcium bridges. PMEs are inhibited by PMEIs in a finely-tuned process (Driouich et al., 2012). Pectins could also be degraded by PLs that are more specifically active on the highly methyl esterified pectins (Yadav et al., 2009). Thus, the high expression of these plant cell wall genes in hydathodes correlates well with the structural specificities of the hydathode cell wall, especially the loose cellular connections observed in the epithem (Cerutti et al., 2017; Cerutti et al., 2019).

#### Inhibitory chemicals and proteins

Several genes involved in flavonoid and lignin synthesis were all less expressed in hydathodes compared to the leaf blade, including the flavonoid activator *MYB75* while the dual flavonoid and lignin repressor *MYB4* was more expressed (Wang et al., 2020) (Supplemental Figure S4). In agreement, phloroglucinol staining revealed lignin deposition in the secondary cell wall in the vasculature but not in the upper part of the hydathode (Supplemental Figure S4). Five genes encoding chitinases which degrade chitin were more expressed inside hydathodes (Table 3). Last, among the most differentially expressed genes, we noted two extensin-related genes (*Pollen Ole 1 like AT3G16670* and *AT3G16660*) and a *defensin-like* (*AT3G05730*) coding for small proteins. These presumably excreted small proteins share very similar gene expression profiles and their expression enhances plant resistance to biotic or oxidative stresses (Ascencio-Ibanez et al., 2008; Luhua et al., 2008) (Table 3).

#### Plant inducible defenses

*ICS1, ICS2, PBS3* and *EDS5*, important genes for SA synthesis and *DRM6*, involved in SA homeostasis, were downregulated in healthy hydathodes (Table 4) (Garcion et al., 2008; Zhang et al., 2017; Rekhter et al., 2019; Torrens-Spence et al., 2019). Accordingly, basal expression of SA-responsive genes *WRKY70*, *PATHOGENESIS-RELATED PROTEIN 2* (*PR2*) and *PHYTOALEXIN-DEFICIENT 4* (*PAD4*) was reduced (Zhou et al., 1998; Li et al., 2004; van Loon et al., 2006). To test for a reduced synthesis of SA inside hydathodes, we quantified SA by mass spectrometry (Figure 6). However, we did not measure any significant difference in SA content. In contrast, hydathodes contained higher concentrations of ABA and less JA compared to the leaf blade (Figure 6). Accordingly, expression of the ABA biosynthetic gene *NCED5* and two ABA-responsive genes was upregulated, whereas expression of the ABA negative regulator *RCK1* was downregulated (Table 4). Expression of *LOX1* and *LOX5* which synthesize the 9(S)-hydroxy-10,12,15-octadecatrienoic acid oxylipin was reduced consistent with the lower JA content and also suggests a reduced accumulation of callose, ROS and defense-related genes upon hydathode infection (Lopez et al., 2011; Yang et al., 2021). Higher expression of *UGT85A1* involved in cytokinin glycosylation is also consistent with the higher content of glycosylated cytokinins measured in hydathode (Jin et al., 2013) whereas a lower expression of *IPT3*, a key determinant of cytokinin biosynthesis in the vasculature was also expected (Nobusawa et al., 2013). Last, genes involved in the synthesis of glucosinolates, which are *Brassicaceae*-specific secondary metabolites promoting plant immunity, were also more expressed in the leaf blade, consistent with a high expression in vascular tissues (Schuster et al., 2006; Redovnikovic et al., 2012) (Supplemental Table S3).

**Figure 6:**
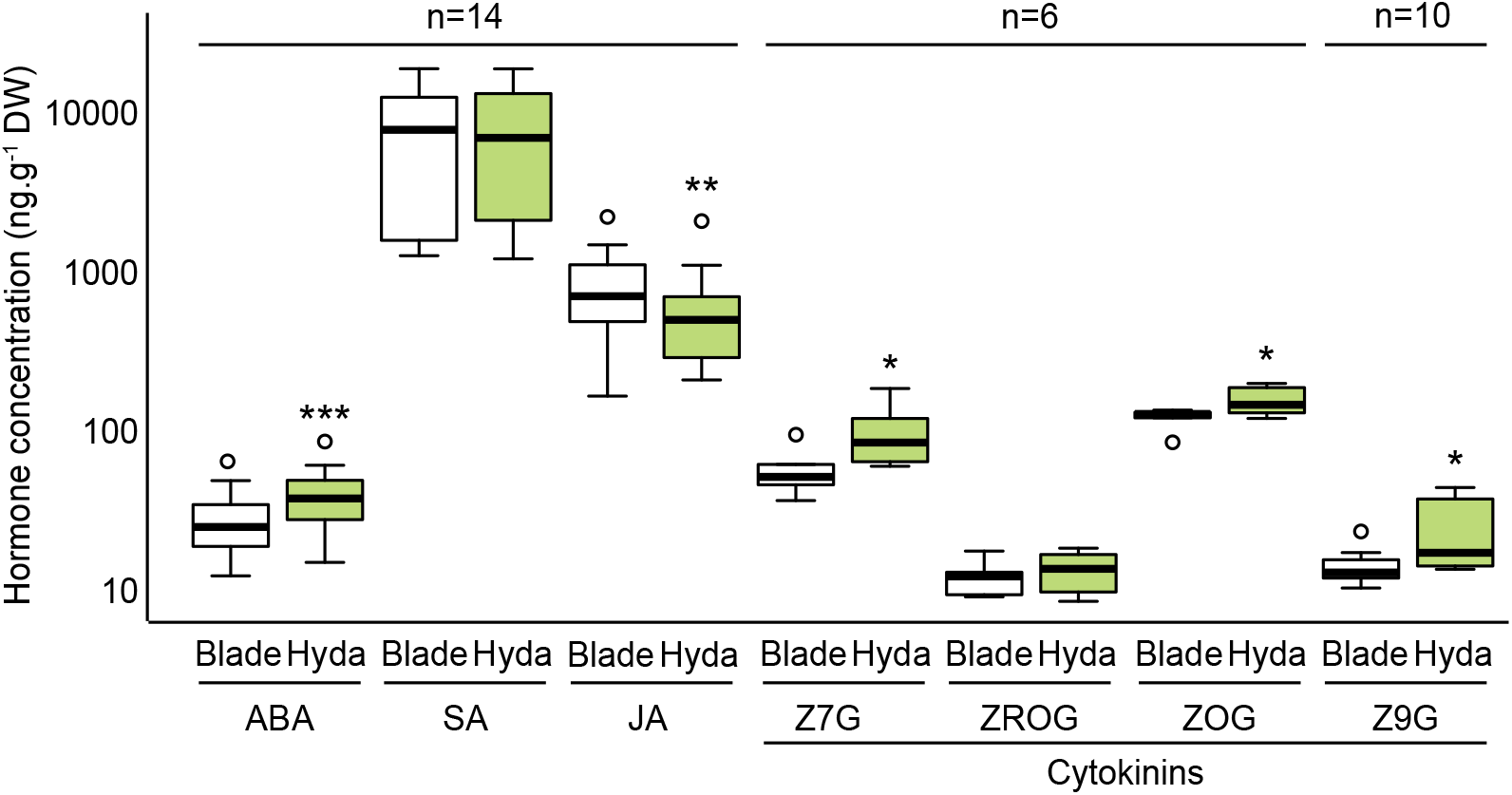
Boxplot representation of plant hormones concentrations in leaf blade (Blade) and hydathodes (Hyda) of eight-week-old plants as measured by GC-MS: abscisic acid (ABA), salicylic acid (SA), jasmonic acid (JA) and cytokinins (zeatin-7-β-D-glucoside (Z7G), trans-zeatin-o-glucoside riboside (ZROG), zeatin-o-glucoside (ZOG), zeatin-9-β-D-glucoside (Z9G). Statistically significant differences were determined using a paired *t*-test (*, p<0.05; ***, p<0.001). n= number of biological replicates.

**Table 4:**
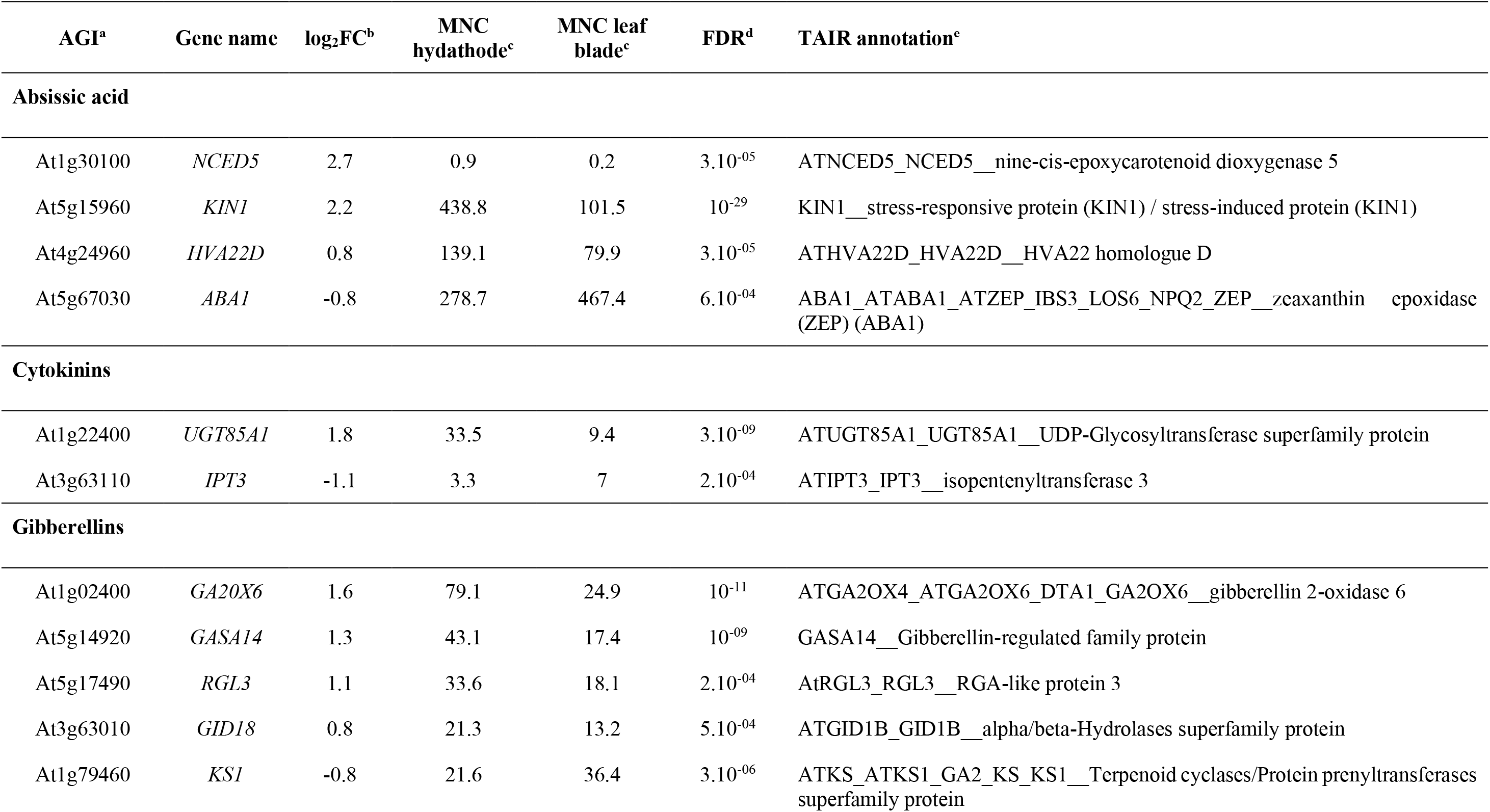
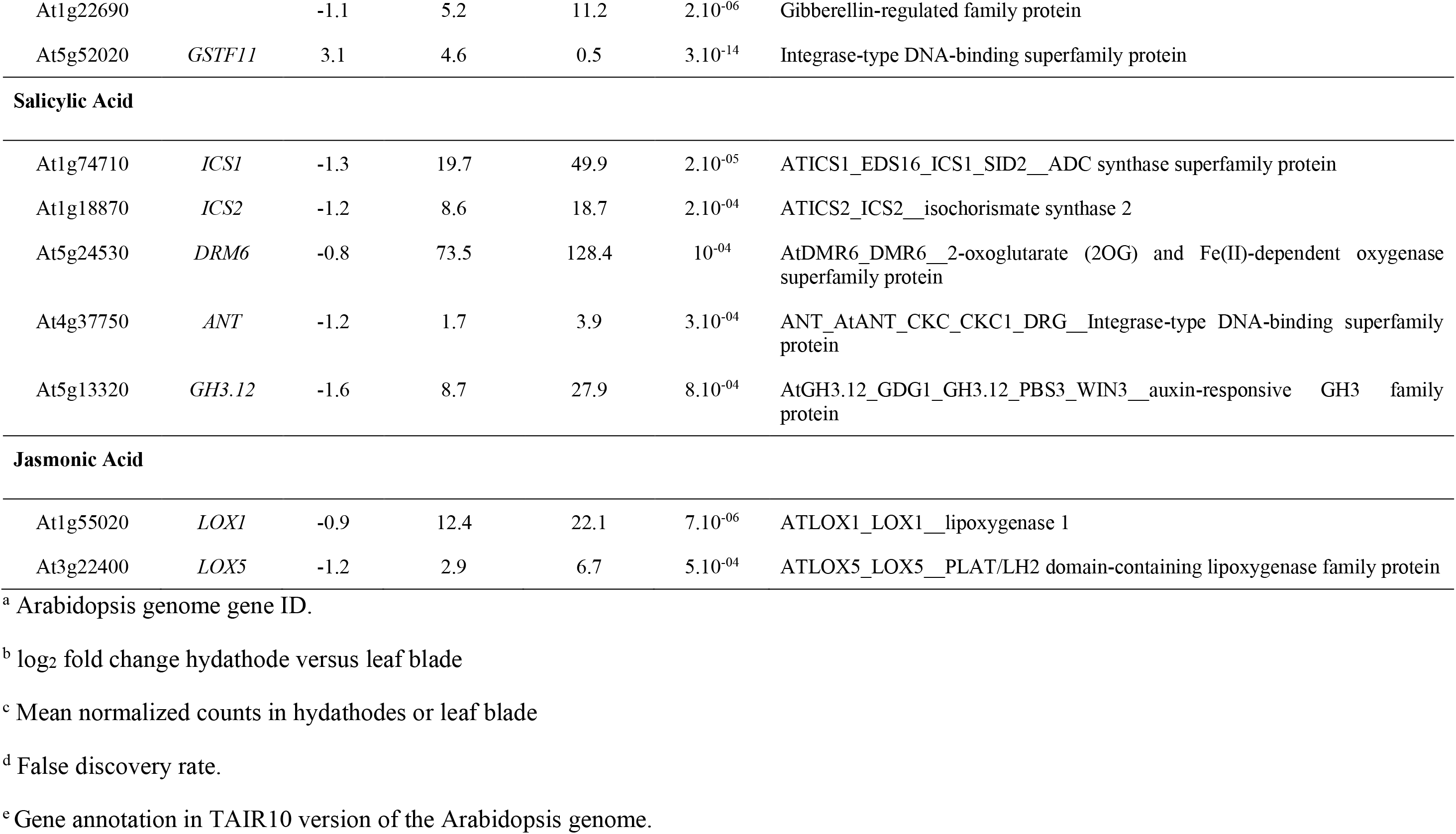
List of the genes involved in hormone metabolism which are differentially expressed in mature Arabidopsis hydathodes relative to the leaf blade (Extracted from Supplemental Table S3, refer to Figure 3 for auxin).

Altogether, basal immunity of healthy hydathodes seems lower than in the leaf blade, especially SA metabolism, and supports the observation that healthy hydathodes may host a rich microbial community (Perrin, 1972). Yet, inducible immune responses upon infection by microbes remain to be determined.

## Discussion

### Hydathodes have a unique transcriptomic identity within the leaf

This analysis identified 1460 genes that are differentially expressed in hydathodes of mature Arabidopsis leaves compared to the neighboring leaf blade tissue and is fully consistent with two recent transcriptomic reports (macrodissected tissues and single cells) which together identified 162 genes preferentially expressed in hydathodes (Kim et al., 2021; Yagi et al., 2021). This work provides a much broader insight into the molecular bases of hydathode physiology and defines a multilayered tissue that is not only structurally, but also physiologically different from neighbor tissues. Because macrodissection yields some contamination by the surrounding mesophyll tissue, both the number of differentially-expressed genes and the intensity of the differential expression are likely underestimated. Only microdissection of hydathode tissues or deeper single cell analyses could further refine the quality of those results. Studying the conservation of those hydathode-specific expression patterns in other plant species would also be of great interest.

### Hydathode transporters prevent the loss of metabolites in the guttation fluid

Significant amounts of xylem sap metabolites reach the hydathodes where many transporters are more expressed. The efficient scavenging of metabolites such as nitrate, Pi, amino acids and sugars could prevent their loss by guttation. We showed that the NRT2.1 and the PHT1;4 high affinity transporters of nitrate and phosphate, respectively, are preferentially expressed in hydathodes (Mudge et al., 2002; Nazoa et al., 2003) and are involved in the active capture of nitrate and Pi in the guttation fluid. In the roots, both NRT2.1 and PHT1;4 are major players for nitrate and Pi uptake from the soil (Filleur et al., 2001; Misson et al., 2004; Shin et al., 2004; Li et al., 2007). Their expression is strongly upregulated by low concentrations of nitrogen (nitrate/ammonium/glutamine) and phosphate, respectively (Nazoa et al., 2003; Misson et al., 2004). Yet, both *NRT2.1* and *PHT1;4* expression in hydathodes is decorrelated from the local nutritional status of the hydathodes as illustrated by the low expression of starvation markers for phosphate (e.g. MGDC, Awai et al., 2001; Kobayashi et al., 2009) and nitrate (NPF7.3/NRT1.5, Cui et al., 2019). Molecular and genetic mechanisms for such hydathode-specific and starvation-independent expression of those transporters remain to be elucidated. For any microbe, the metabolites that could be present in the guttation fluid could represent valuable nutrients. In the bacterial pathogen *Xanthomonas campestris* pv. *campestris*, the causal agent of back rot disease in *Brassicaceae*, the expression of transporters of inorganic phosphate (Pi), sulfur or nitrogen was upregulated inside cauliflower hydathodes (Luneau et al., 2022) and Pi uptake limits growth inside hydathodes (Luneau et al., 2022; Luneau et al., 2022). This suggests that active scavenging of nutrients in hydathodes could limit proliferation of microbes and contribute to the immunity of this organ, otherwise fully exposed to the epiphytic microbes.

### Hydathodes produce and accumulate auxin and its conjugates

Direct hormone quantification showed that both free auxin and oxindole-3-AIA, a down-product of the auxin storage and inactivation pathway (Hayashi et al., 2021) accumulate in hydathodes. During leaf margin differentiation, such auxin maxima promote serration outgrowth through cell proliferation and patterning (Kawamura et al., 2010; Kasprzewska et al., 2015). Auxin also plays a role during xylem specification and differentiation and may explain the very dense atypical delta shape structure of the xylem irrigating the epithem (Aloni, 2013; Kondo et al., 2014). More than 67 genes involved in auxin biosynthesis, regulation, signaling, conjugation and response are upregulated in hydathodes thus confirming and extending our knowledge about the importance of auxin metabolism in hydathode biology. This includes several *YUCCA* genes that encode flavin monooxygenase-like enzymes that catalyze the rate-limiting step in Trp-dependent auxin biosynthesis and *TAA1* which performs the first committed steps of auxin synthesis. In contrast, expression of several auxin-related transporters genes was down-regulated. These observations are in agreement with the hypothesis of a higher accumulation of auxin in hydathodes (Aloni et al., 2003). This is also supported by the overexpression in hydathode of numerous copies of auxin biosynthesis regulators belonging to the STYLISH, NGA or TCP families or differential expression of many auxins signaling genes belonging to the AUX-IAA, ARF and SAUR families. Yet, the exact biological functions of such a high auxin metabolism in fully differentiated and mature hydathodes remains enigmatic.

These transcriptomic and physiological results provide molecular insights on how hydathodes actively scavenge xylem sap-derived metabolites and establish a robust foundation for future hydathode research.

## Materials and methods

### Plant material and growth conditions

Arabidopsis (Accession Col-0 or Ws-4 and mutant lines) was grown in a growth chamber on *Jiffy-7*® peat pellets (http://www.jiffypot.com) under short-day conditions (8h light, 100-120μE). Promoter:*GUS* reporter lines were obtained from NASC stock center: *TAA1*, N66987; *YUC2*, N69892; *YUC5*, 69894; *YUC8*, N69897; *YUC9*, N69945. DR5-GUS and *pht1;4-1* mutant (Ws-4 ecotype; (Misson et al., 2004)) and *pGDU4-1-GUS* reporter line from Guillaume Pilot (Virginia Tech, Blacksburg, VA 24061, USA). The *DR5:VENUS* line was described in Maugarny Cales *et al*. (2019) and the E325-GFP enhancer line used as a hydathode marker line (Yagi et al., 2021) was obtained from the NASC stock center.

### Tissue sampling for transcriptomic

Hydathodes were harvested on nine- to ten-week-old Arabidopsis plants. Teeth containing the hydathode on the distal part of intermediate and/or fully extended leaves and leaf blade samples (comprising mesophyll, stomata, epiderm and some minor veins) were excised with a razor blade. Samples were directly frozen in liquid nitrogen and stored at −80°C until RNA extraction. The sampling was performed for four biological replicates over a period of one year.

### RNA extraction, sequencing and differential expression analyses

RNA extraction was performed as described (Chomczynski, 1993). Global gene expression profiles were determined by replicated strand-specific Illumina RNA-Seq. Paired-end oriented RNA sequencing (2×150 pb) was performed on an Illumina HiSeq3000 using the Illumina HiSeq3000 Reagent Kits. Paired-end reads were mapped on the reference Arabidopsis genome using the glint software (release 1.0.rc12.826:833, http://lipm-bioinfo.toulouse.inra.fr/download/glint). Only best-scores were taken into account, with a minimal hit length of 60 bp, a maximum of 5 mismatches and no gap allowed. Ambiguous matches with the same best score were removed (parameters: glint mappe --mate-maxdist 10000 -C 0 --output-format bed --best-score --no-gap --step 2 --lmin 60 --mmis 5).

Differential expression analysis was performed with R (v3.6.2) using the EdgeR package (v3.28.1) and normalization with the TMM method (trimmed mean of M-values, Robinson and Oshlack, 2010) as described (Pecrix et al., 2018). Genes with no count across all libraries were not retained for further analysis. Differentially-expressed genes (DEGs) were called using the GLM likelihood ratio test, with a FDR adjusted q-value < 0.001. For quality control, plots of normalized datasets, heatmap on sample-to-sample Euclidean distances were generated with the package pheatmap version 1.0.8. Enrichment analysis used the MAPMAN software (http://bar.utoronto.ca/ntools/cgi-bin/ntools_classification_superviewer.cgi).

### Statistical Analyses

All indications about statistical analyses are indicated in Figure legends.

### Visualization of ß-glucuronidase activity *in planta*

Plant tissues were vacuum-infiltrated in 50 mM sodium phosphate buffer, pH 7.2, 1 mM 5-bromo-4-chloro-3-indolyl β-D-glucuronide, 0.2% Triton X-100 [v/v], 2 mM potassium ferricyanide [K_3_FeCN_6_], 2 mM potassium ferrocyanide [K_4_FeCN_6_]) and incubated from 2 to 12 hours at 37°C. Samples were clarified by successive incubations of 80% ethanol at 60°C for few hours and maintained in same solution at room temperature for several days.

### Confocal imaging

Leaves from eight-week-old plants grown in short day condition were fixed in 4% paraformaldehyde under a vacuum for 1h and cleared in Clearsee (10% xylitol, 25% urea, 15% deoxycholate) (Kurihara et al., 2015) and calcofluor (0.1%). Confocal imaging was performed on a Leica SP5 inverted microscope (Leica Microsystems, Wetzlar, Germany) using a Leica 40x HCX PL APO CS lens. Calcofluor was excited at 405nm and visualized between 410-450nm. GFP and YFP were excited at 488nm and 514nm and visualized between 518-554nm and 518-543 nm respectively. Max intensity projections along the Z axis were made using ImageJ and FigureJ (Mutterer and Zinck, 2013) was used to assemble the figure.

### Binary vectors and plant transformation

For promoter expression analyses, ca. 2-kb promoter region was amplified by PCR from *Arabidopsis* Col-0 genomic DNA with Gateway-compatible oligonucleotides (Supplementary Table S5), cloned into the Gateway donor vector pDON207 by BP reaction (Invitrogen) and sequenced. Resulting ENTRY clones were recombined by LR recombination into pKGWFS7 binary vector (Karimi et al., 2002) yielding a construct with the promoter driving the transcription of the GFP:GUS translational fusion. Plants were transformed as described (Chung et al., 2000).

### Metabolomic analyses

Samples were collected on four-week-old plants. The day before sample harvest, at the end of the light period, for each fully developed leaf, the 0.5 to 1mm leaf margin was removed with a scissor on one half of the leaf when the other half was left intact. The plants were watered and covered to keep them at 100% relative humidity. Guttation droplets and pre-hydathode xylem sap were harvested at the end of the night period from intact and excised leaf margins, respectively. Xylem fluid was then collected on petiole after leaf removal after 3h in the dark and 100% relative humidity. About 500, 100 and 100μl of lyophilized fluid were used for metabolome analysis of guttation fluid, pre-hydathode xylem sap and xylem sap before leaf, respectively. These samples were extracted, derivatized and analyzed by an Agilent 7890A gas chromatograph coupled to an Agilent 5975C mass spectrometer as described (Fiehn, 2006; Masclaux-Daubresse et al., 2014). Data were analyzed with AMDIS (http://chemdata.nist.gov/mass-spc/amdis/) and QuanLynx software (Waters).

Colorimetric nitrate and inorganic phosphate (molybdenum blue method) contents were determined as described (Kanno et al., 2016; Zhao and Wang, 2017).

For hormone measurements, each sample of hydathodes and leaf blade were harvested with a 1.5mm Miltex® biopsy Punch. Six mg of dry powder were extracted with 0.8 mL of acetone/water/acetic acid (80/19/1 v:v:v). Abscisic acid, salicylic acid, jasmonic acid, indole-3-acetic acid and cytokinins stable labeled isotopes used as internal standards were prepared as described (Le Roux et al., 2014). One ng of each and 0.5ng cytokinins standard was added to the sample. The extract was vigorously shaken for 1min, sonicated for 1 min at 25 Hz, shaken for 10 minutes at 10°C in a Thermomixer (Eppendorf®) and then centrifuged (8,000g, 10 °C, 10 min.). The supernatants were collected and the pellets were re-extracted twice with 0.4 mL of the same extraction solution, vigorously shaken (1 min) and sonicated (1 min; 25 Hz). After the centrifugations, the three supernatants were pooled and dried (Final Volume 1.6 mL). Each dry extract was dissolved in 100 μL of acetonitrile/water (50/50 v/v), filtered and analyzed using a Waters Acquity ultra performance liquid chromatography coupled to a Waters Xevo Triple quadrupole mass spectrometer TQS (UPLC-ESI-MS/MS) as described (Le Roux et al., 2014).

## Acknowledgements

This work was supported by a grant from the Agence Nationale de la Recherche NEPHRON project (ANR-18-CE20-0020-01) to JMR, CB, GC, SCI, CR, MC, LF, SCH, DV, MFJ, SCA, LNU, PL, NLE, LNA and LDN. This study is set within the framework of the ‘Laboratories d’Excellences’ (LABEX) TULIP (ANR-10-LABX-41) and of the ‘Ecole Universitaire de Recherche’ (EUR) TULIP-GS (ANR-18-EURE-0019). The IJPB benefits from the support of Saclay Plant Sciences-SPS (ANR-17-EUR-0007). This work has benefited from the support of IJPB’s Plant Observatory technological platforms. We thank Guillaume Pilot for the gift of the *pGDU4-1:GUS* reporter line (Virginia Tech, Blacksburg, VA 24061, USA).

## Author contributions

JMR, LNU, PL, NLE, LNA, MS and LDN conceived the study and supervised experiments. JMR, CB, GC, SCI, CR, MC, LF, DV, SCH and NLE performed and analyzed the experiments. CB, MFJ, SCA and JMR processed and analyzed transcriptomic data; JMR and LDN drafted the manuscript. All authors critically reviewed the results and the manuscript.

## Data availability statement

All relevant data are within the manuscript and its Supporting Information files. RNA sequencing results are available in the single read archive SRP322413 (SRX11051786, SRX11051788-9 and SRX11051791-5).

## Supplemental Figures and Tables

**Table S1:** Comparison of gene expression in hydathodes *vs* leaf blade obtained from promoter:*GUS* fusions and this transcriptomic analysis

**Table S2:** RNA sequencing results for libraries made from hydathodes and leaf blade tissues sampled on mature leaves of Arabidopsis accession Col-0.

**Table S3:** Expression data and properties of all genes in hydathodes versus leaf blade in mature Arabidopsis leaves from accession Col-0.

**Table S4:** Concentration of 52 metabolites in fluids collected at the petiole (P), before hydathodes (BH) or at the margin (guttation fluid, GF) of leaves from four-week-old Arabidopsis plants were measured with GC-MS.

**Table S5:** Name and sequence of oligonucleotides used in this study.

**Figure S1:** Transcriptomic analysis of Arabidopsis hydathode-enriched versus leaf blade samples (A) Analysis of euclidean distances between global expression profiles in the four biological replicates per condition cluster samples per tissue. A color key is shown on the right. (B) Volcano plot of gene expression levels (log_2_ fold change (FC)) in hydathodes versus leaf blade and false discovery rate (log_10_FDR). Genes with an FDR below 0.001 and absolute value (log_2_FC) higher than 1 are indicated in blue and red when overexpressed in the hydathodes and leaf blade samples, respectively. (C) Functional categories significantly over- or underrepresented in hydathodes (red) and leaf blade (blue) using a MapMan classification. Analysis was applied to the annotated DEG (FDR<0.001). Normed frequency number indicates the proportion of DEG in the functional category (raw numbers are indicated; http://bar.utoronto.ca/ntools/cgi-bin/ntools_classification_superviewer.cgi).

**Figure S2:** Visualization of promoter activities in leaves of four- and eight-week-old transgenic Arabidopsis plants carrying promoter: *GUS* reporter fusions. *TAA1* and *YUCs* genes are involved in auxin synthesis. The *DR5* synthetic promoter is a reporter of auxin response. *PHT1;4* (studied using a promoter trap mutant) and *GDU4.7* encode phosphate and glutamate transporters, respectively. Expression fold change (log_2_FC) between hydathodes versus leaf blade is indicated as described (Table S3). Scale bar= 200μm.

**Figure S3:** Observation of the most distal or fourth most distal hydath-ode on a mature leaf of an eight-week-old plant by confocal microscopy in the *DR5:VENUS* auxin signaling reporter line (Yellow) and the *E325:GFP* enhancer trap line (Green). Bright field and bright field+ fluorescence overlay images are shown. Scale bars= 100μm.

**Figure S4:** Genes involved in phenylpropanoid metabolism are relatively less expressed in mature hydathodes relative to the leaf blade. (A) DEG involved in the biosynthesis of lignins and flavonoids and their regulators. Numbers indicate their log_2_ fold change as described (Table S3). (B) Lignin staining of a Arabidopsis hydathode from an eight-week-old plant using phloroglucinol. Lignified secondary cell wall of xylem vessels (visualized in pink with phloroglucinol staining) is restricted to the lower portion of hydathodes and the vascular elements. Scale bar= 200μm.

