## Supplemental figures for "Arabidopsis hydathodes are sites of intense auxin metabolism and nutrient scavenging"

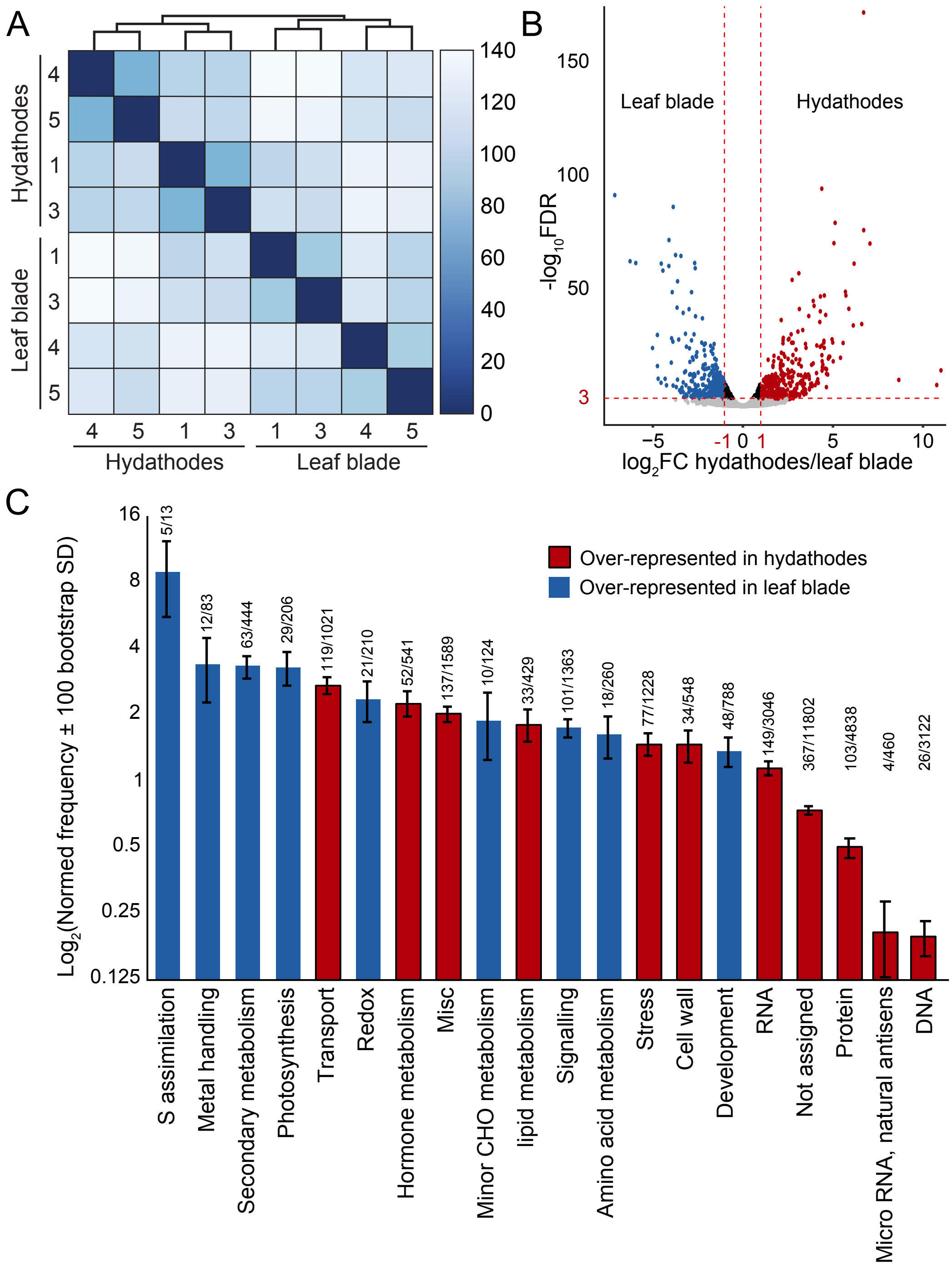


**Figure S1:** Transcriptomic analysis of Arabidopsis hydathode-enriched versus leaf blade samples (A) Analysis of euclidean distances between global expression profiles in the four biological replicates per condition cluster samples per tissue. A color key is shown on the right. (B) Volcano plot of gene expression levels ($log_2$ fold change (FC)) in hydathodes versus leaf blade and false discovery rate ($log_{10}$FDR). Genes with an FDR below 0.001 and absolute value ($log_2$FC) higher than 1 are indicated in blue and red when overexpressed in the hydathodes and leaf blade samples, respectively. (C) Functional categories significantly over- or under-represented in hydathodes (red) and leaf blade (blue) using a MapMan classification. Analysis was applied to the annotated DEG (FDR<0.001). Normed frequency number indicates the proportion of DEG in the functional category (raw numbers are indicated; http://bar.utoronto.ca/ntools/cgi-bin/ntools_classification_superviewer.cgi).

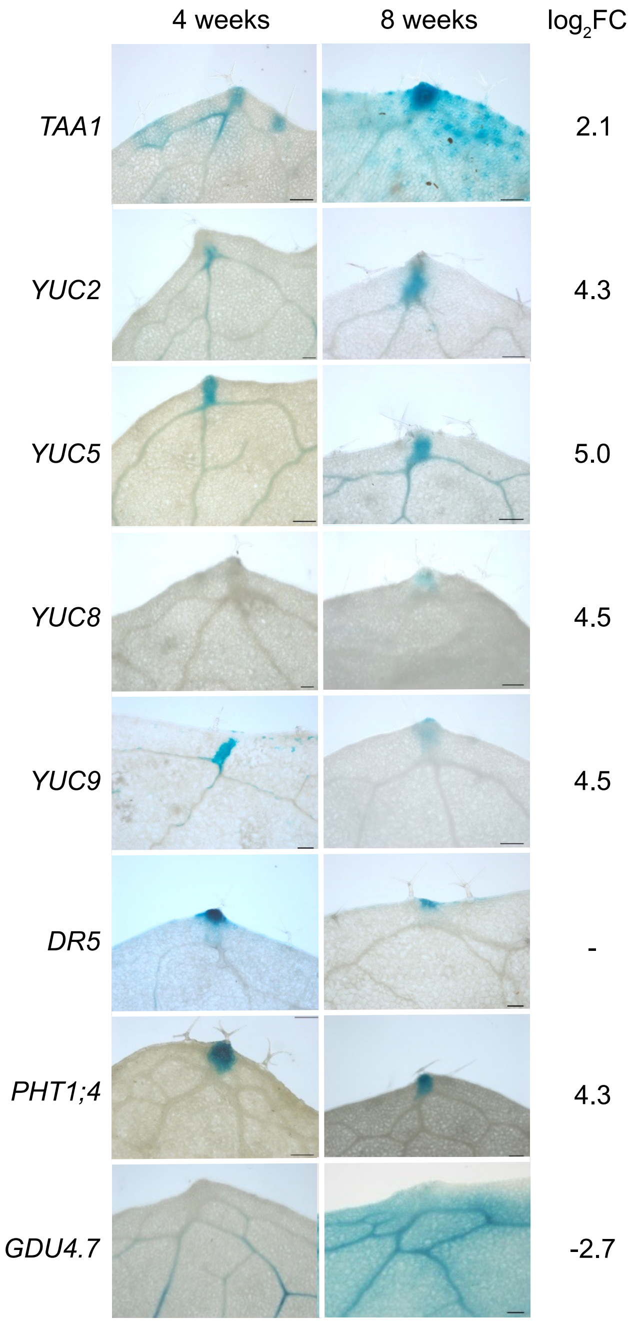


**Figure S2:** Visualization of promoter activities in leaves of four- and eight-week-old transgenic Arabidopsis plants carrying promoter:*GUS* reporter fusions. *TAA1* and *YUCs* genes are involved in auxin synthesis. The *DR5* synthetic promoter is a reporter of auxin response. *PHT1;4* (studied using a promoter trap mutant) and *GDU4.7* encode phosphate and glutamate transporters, respectively. Expression fold change ($log_2FC$) between hydathodes versus leaf blade is indicated as described (Supplemental Table S3). Scale bar= 200µm.

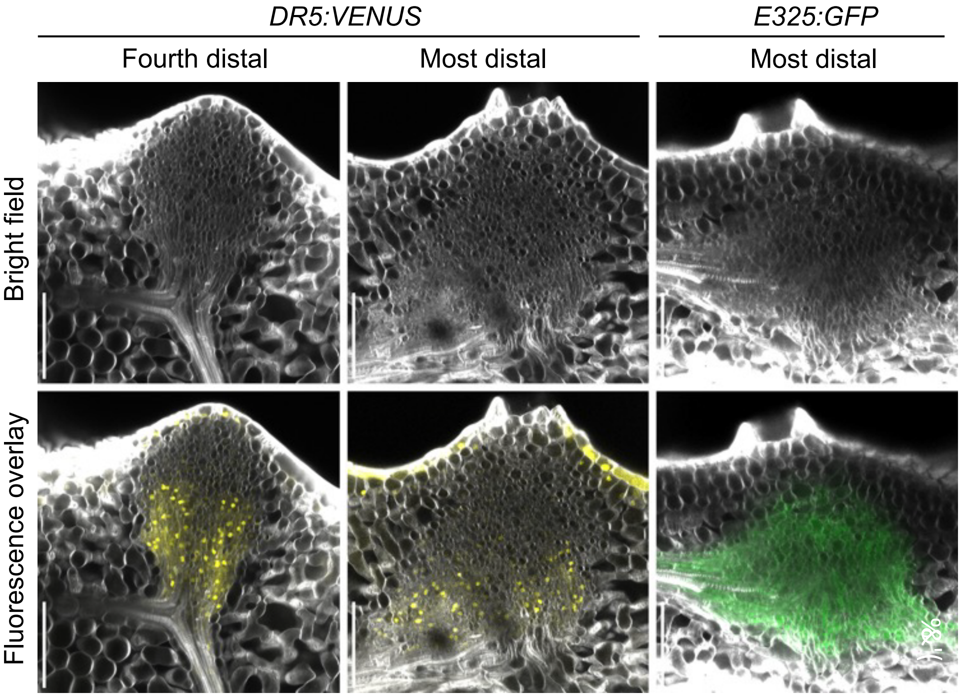


**Figure S3:** Observation of the most distal or fourth most distal hydathode on a mature leaf of an eight-week-old plant by confocal microscopy in the *DR5:VENUS* auxin signaling reporter line (Yellow) and the *E325:GFP* enhancer trap line (Green). Bright field and bright field+ fluorescence overlay images are shown. Scale bars= 100µm.

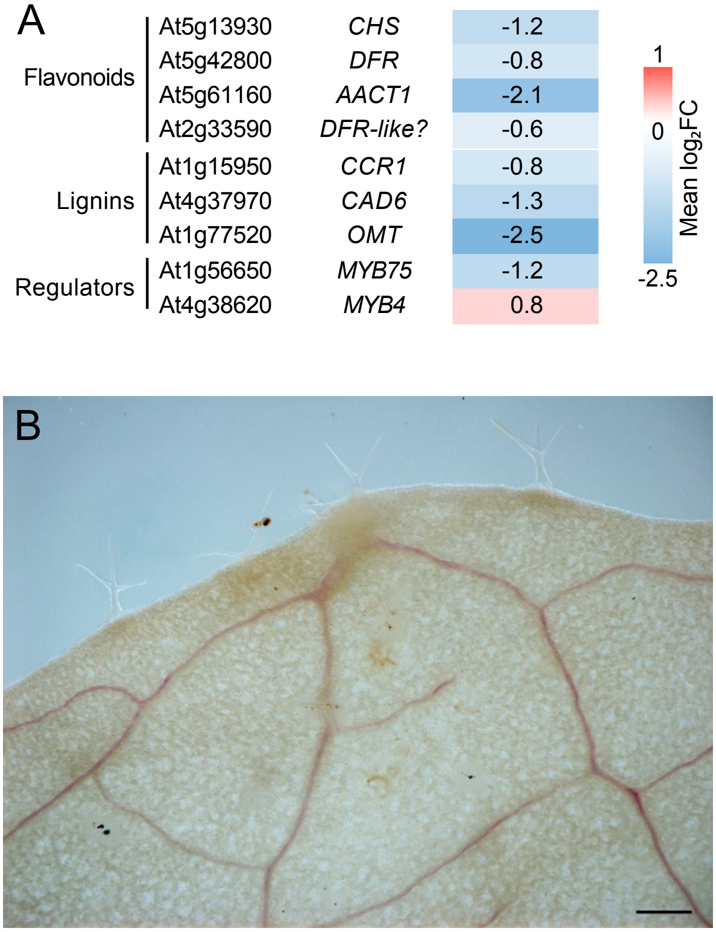


**Figure S4:** Genes involved in phenylpropanoid metabolism are relatively less expressed in mature hydathodes relative to the leaf blade. (A) DEG involved in the biosynthesis of lignins and flavonoids and their regulators. Numbers indicate their $\log_2$ fold change ($\log_2$FC) as described (Supplemental Table S3). (B) Lignin staining of a Arabidopsis hydathode from an eight-week-old plant using phloroglucinol. Lignified secondary cell wall of xylem vessels (visualized in pink with phloroglucinol staining) is restricted to the lower portion of hydathodes and the vascular elements. Scale bar= 200µm.
