## Supplemental Tables for "Arabidopsis hydathodes are sites of intense auxin metabolism and nutrient scavenging"

**Table S1:** Comparison of gene expression in hydathodes *vs* leaf blade obtained from promoter:*GUS* fusions and this transcriptomic analysis

| **AGI[a]** | **Gene name** | **$Log_2FC$[b]** | **FDR[c]** | **References** |
|---|---|---|---|---|
| **Genes preferentially expressed in hydathodes based on promoter:*GUS* fusions analyses** | | | | |
| At1g01030 | *NGA3* | 3.1 | $3.10^{-19}$ | (Alvarez et al., 2009; Trigueros et al., 2009) |
| At1g08090 | *NRT2.1* | 6.6 | $10^{-36}$ | (Nazoa et al., 2003) |
| At1g09970 | *RLK7* | 0.3 | 0.54 | (Wu et al., 2016) |
| At1g25320 | | 0.7 | $7.10^{-3}$ | (Wu et al., 2016) |
| At1g28230 | *PUP1* | 1.2 | $3.10^{-6}$ | (Burkle et al., 2003) |
| At1g30350 | *PLL7* | - | - | (Sun and van Nocker, 2010) |
| At1g30450 | *ATCCC* | 0.0 | 0.9 | (Colmenero-Flores et al., 2007) |
| At1g66830 | | 2.5 | $6.10^{-30}$ | (Wu et al., 2016) |
| At1g75520 | *SRS5* | 2.8 | $2.10^{-4}$ | (Baylis et al., 2013) |

| | | | | |
|---|---|---|---|---|
| At1g75640 | | 0.2 | 0.9 | (Wu et al., 2016) |
| At1g78000 | *SULTR1.2* | 2.7 | $10^{-6}$ | (Shibagaki et al., 2002) |
| At2g01210 | | 0.2 | 0.9 | (Wu et al., 2016) |
| At2g16120 | *PMT1* | - | - | (Klepek et al., 2010) |
| At2g26650 | *AKT1* | 1.1 | $10^{-7}$ | (Lagarde et al., 1996; Colmenero-Flores et al., 2007) |
| At2g28970 | | 6.1 | $4.10^{-36}$ | (Wu et al., 2016) |
| At2g38940 | *PHT1;4* | 4.3 | $7.10^{-49}$ | (Misson et al., 2004) |
| At3g06120 | *MUTE* | 0.9 | 0.4 | (Pillitteri et al., 2008) |
| At3g09220 | *LAC7* | 0.7 | 0.7 | (Turlapati et al., 2011) |
| At3g27400 | *PLL18* | 1.0 | $2.10^{-2}$ | (Sun and van Nocker, 2010) |
| At3g29160 | *SnRK1.2* | -0.1 | 0.9 | (Williams et al., 2014) |
| At3g51060 | *STYLISH1* | 5.9 | $2.10^{-43}$ | (Baylis et al., 2013) |
| At3g54420 | *EP3* | 1.6 | $4.10^{-14}$ | (Passarinho et al., 2001) |
| At3g55140 | *PLL2* | -0.2 | 0.4 | (Sun and van Nocker, 2010) |
| At3g57480 | *SAP13* | -0.1 | 0.8 | (Dixit et al., 2018) |
| At4g13260 | *YUC2* | 4.3 | $10^{-37}$ | (Muller-Moule et al., 2016) |
| At4g13280 | *TPS12* | - | - | (Ro et al., 2006) |
| At4g13300 | *TPS13* | - | - | (Ro et al., 2006) |

| | | | | |
|---|---|---|---|---|
| At4g28270 | *RMA2* | 0.1 | 0.9 | (Son et al., 2009) |
| At4g32650 | *KC1* | 2.8 | $2.10^{-28}$ | (Pilot et al., 2003) |
| At4g35020 | *ROP6* | 0.1 | 0.9 | (Poraty-Gavra et al., 2013) |
| AT4G36260 | *STY2* | 6,2 | $2.10^{-63}$ | (Baylis et al., 2013) |
| At4g37870 | *PEPCK* | 1.8 | $10^{-14}$ | (Penfield et al., 2012) |
| At4g37990 | *CAD8* | 1.5 | $5.10^{-4}$ | (Kim et al., 2007) |
| At5g08150 | *SOB5* | 0.1 | 0.9 | (Zhang et al., 2006) |
| At5g11320 | *YUC4* | 5.0 | $2.10^{-20}$ | (Wang et al., 2011) |
| At5g28030 | *DES1* | 3.1 | $2.10^{-43}$ | (Laureano-Marin et al., 2014) |
| At5g43350 | *PHT1;1* | -0.9 | 0.5 | (Mudge et al., 2002) |
| At5g46330 | *FLS2* | 0.4 | $3.10^{-2}$ | (Beck et al., 2014) |
| At5g53320 | | 3.4 | $2.10^{-13}$ | (Wu et al., 2016) |
| At5g66350 | *SHI* | 3,3 | $4.10^{-34}$ | (Baylis et al., 2013) |
| At5g63180 | *PLL15* | 0.2 | 0.8 | (Sun and van Nocker, 2010) |
| **Genes expressed in both hydathodes and leaf blade based on promoter:*GUS* fusions analyses** | | | | |
| At1g01130 | *ISA2* | 0.4 | 0.8 | (Li et al., 2007) |
| At1g10130 | *ECA3* | 0.0 | 0.9 | (Mills et al., 2008) |
| At1g26730 | *PHO1;H7* | 0.4 | 0.4 | (Wang et al., 2004) |

| | | | | |
|---|---|---|---|---|
| At1g41830 | *SKS6* | -0.5 | $2.10^{-2}$ | (Jacobs and Roe, 2005) |
| At1g56120 | | 0.2 | 0.8 | (Wu et al., 2016) |
| At1g59870 | *PDR8* | -0.5 | 0.2 | (Kobae et al., 2006) |
| At2g02860 | *SUT2* | 0.0 | 1 | (Schulze et al., 2000) |
| At2g16130 | *PMT2* | - | - | (Klepek et al., 2010) |
| At2g37330 | *ALS3* | 0.6 | $10^{-2}$ | (Larsen et al., 2005) |
| At2g47585 | *mir164A* | - | - | (Bazzini et al., 2009) |
| At3g07930 | *MBD4L* | -0.1 | 0.9 | (Nota et al., 2015) |
| At3g09540 | *PLL1* | 0.4 | $3.10^{-2}$ | (Sun and van Nocker, 2010) |
| At3g19450 | *CAD4* | -0.3 | 0.4 | (Kim et al., 2007) |
| At3g24230 | *PLL24* | 0.2 | 0.9 | (Sun and van Nocker, 2010) |
| At3g24670 | *PLL22* | 0.1 | 0.9 | (Sun and van Nocker, 2010) |
| At3g25230 | *ROF1* | 0.1 | 0.9 | (Aviezer-Hagai et al., 2007) |
| At4g02570 | *AXR6* | 0.1 | 0.9 | (Esteve-Bruna et al., 2013) |
| At4g09020 | *ISA3* | 0.1 | 0.8 | (Li et al., 2007) |
| At4g27080 | *PDI7* | 0.1 | 0.6 | (Yuen et al., 2016) |
| At4g34230 | *CAD5* | 0.7 | $3.10^{-2}$ | (Kim et al., 2007) |
| At4g37970 | *CAD6* | -1.3 | $4.10^{-5}$ | (Kim et al., 2007) |

| | | | | |
|---|---|---|---|---|
| At4g37980 | *CAD7* | 1.7 | $2.10^{-20}$ | (Kim et al., 2007) |
| At5g14800 | *P5R* | 0.5 | $10^{-2}$ | (Hua et al., 1997) |
| At5g20270 | *HHP1* | 0.1 | 0.8 | (Chen et al., 2010) |
| At5g26340 | *STP13* | 0.1 | 0.8 | (Schofield et al., 2009) |
| At5g33280 | *CLCg* | -0.3 | 0.2 | (Nguyen et al., 2016) |
| At5g43360 | *PHT1;3* | - | - | (Mudge et al., 2002) |
| At5g43890 | *YUC5* | 5.0 | $5.10^{-30}$ | (Muller-Moule et al., 2016) |
| **Genes preferentially expressed in leaf blade based on promoter:*GUS* fusions analyses** | | | | |
| At5g37770 | *CML24* | -0.1 | 0.8 | (Delk et al., 2005) |
| At2g24762 | *GDU4* | -2,7 | $10^{-14}$ | (Pratelli et al., 2010) |
| At2g39450 | *MTP11* | -0.1 | 0.9 | (Peiter et al., 2007) |
| At2g46340 | *SPA1* | -0.6 | $3.10^{-4}$ | (Ranjan et al., 2011) |
| At3g17690 | *CNGC19* | -0.4 | 0.6 | (Kugler et al., 2009) |
| At3g18440 | *ALMT9* | 0.2 | 0.5 | (Kovermann et al., 2007) |
| At3g21670 | *NPF6.4* | -1.0 | $10^{-3}$ | (Tong et al., 2016) |
| At3g51920 | *CML9* | -0.9 | $3.10^{-04}$ | (Magnan et al., 2008) |
| At4g31730 | *GDU1* | -2.1 | $2.10^{-02}$ | (Pratelli et al., 2010) |
| At3g19710 | *BCAT4* | -4.4 | $3.10^{-60}$ | (Schuster et al., 2006) |

| | | | | |
|---|---|---|---|---|
| At5g61420 | *MYB28* | -0.9 | $5.10^{-8}$ | (Gigolashvili et al., 2007) |
| At5g23010 | *MAM1* | -3.7 | $5.10^{-44}$ | (Redovnikovic et al., 2012) |

[a] Arabidopsis genome gene ID.

[b] $Log_2$ fold change hydathode versus leaf blade (This study, Supplemental Table S3)

[c] False discovery rate (This study, Supplemental Table S3)

**Table S2:** RNA sequencing results for libraries made from hydathodes and leaf blade tissues sampled on mature leaves of Arabidopsis accession Col-0.

| Sample | Biological replicates | Library name | Number of raw reads | Uniquely mapped reads (%)[a] | SRA accession |
|---|---|---|---|---|---|
| **Hydathode** | 4 | ARATH-9-10W-ShortDays-Hydathode-C1 | 22 328 928 | 94.1 | SRX11051786 |
| | | ARATH-9-10W-ShortDays-Hydathode-C3 | 32 660 149 | 94.6 | SRX11051788 |
| | | ARATH-9-10W-ShortDays-Hydathode-C4 | 14 503 424 | 89.7 | SRX11051794 |
| | | ARATH-9-10W-ShortDays-Hydathode-C5 | 10 811 758 | 83.2 | SRX11051795 |
| **Mesophyll** | 4 | ARATH-9-10W-ShortDays-Mesophyll-C1 | 28 890 042 | 94.3 | SRX11051789 |
| | | ARATH-9-10W-ShortDays-Mesophyll-C3 | 35 632 940 | 94.1 | SRX11051791 |
| | | ARATH-9-10W-ShortDays-Mesophyll-C4 | 11 877 761 | 88.8 | SRX11051792 |
| | | ARATH-9-10W-ShortDays-Mesophyll-C5 | 12 500 529 | 90.3 | SRX11051793 |

[a] Percentage of raw reads mapped uniquely to the annotated sequences of *Arabidopsis thaliana* chromosomes (TAIR10).
[b] Sequence Read Archive accession numbers (SRP322413)

**Table S3:** Expression data and properties of all genes in hydathodes versus leaf blade in mature Arabidopsis leaves from accession Col-0.

**Table S4:** Concentration of 52 metabolites in fluids collected at the petiole (P), before hydathodes (BH) or at the margin (guttation fluid, GF) of leaves from four-week-old Arabidopsis plants were measured with GC-MS.

| **Compound category** | **Compound name** | **Mean concentration (nmol/mL)** [a] | | | **Median concentration (nmol/mL)** [a] | | | **Kluskal-Wallis test ($p<0.05$)** [b] | | |
|---|---|---|---|---|---|---|---|---|---|---|
| | | Fluid at petiole level | Pre-hydathode fluid | Guttation fluid | Fluid at petiole level | Pre-hydathode fluid | Guttation fluid | Petiole vs pre-hydathode | Petiole vs guttation fluid | Pre-hydathode vs guttation fluid |
| **Amino acids** | **GABA** [c] | 140.7 | 5.59 | 0.56 | 88.61 | 4.64 | 0.22 | a | b | c |
| | **Glutamate** | 25.86 | 1.92 | 0.37 | 10.78 | 1.32 | 0.13 | a | b | c |
| | **Aspartate** | 38.5 | 3.52 | 0.36 | 7.91 | 0.32 | 0.045 | a | b | c |
| | **Phenylalanine** | 5.91 | 0.56 | 0.17 | 5.36 | 0.38 | 0.089 | a | b | c |
| | **Lysine** | 8.39 | 0.46 | 0.16 | 9.44 | 0.24 | 0.068 | a | b | c |
| | **Alanine** | 30.33 | 2.01 | 0.78 | 23.72 | 1.04 | 0.30 | a | b | c |
| | **Threonine** | 13.5 | 1.36 | 0.36 | 14.34 | 0.69 | 0.19 | a | b | c |
| | **Valine** | 27.37 | 1.62 | 0.63 | 24.32 | 0.97 | 0.26 | a | b | c |

| | | | | | | | | | |
|---|---|---|---|---|---|---|---|---|---|
| **Leucine** | 12.82 | 1.05 | 0.45 | 10.85 | 0.62 | 0.19 | a | b | c |
| **Isoleucine** | 13.13 | 0.72 | 0.33 | 14.25 | 0.46 | 0.14 | a | b | c |
| **Tyrosine** | 3.87 | 1.7 | 0.23 | 4.22 | 0.18 | 0.055 | a | b | c |
| **Serine** | 17.33 | 2.98 | 0.79 | 16.40 | 1.06 | 0.43 | a | b | c |
| **Glycine** | 8.44 | 2.38 | 1.01 | 6.86 | 1.05 | 0.55 | a | b | c |
| Methionine | 1.42 | 0.06 | 0.01 | 0.99 | 0.0087 | 0.0034 | a | b | b |
| Tryptophan | 0.32 | 0.02 | 0.01 | 0.11 | 0.0098 | 0.0008 | a | b | b |
| Glutamine | 1497.08 | 43.72 | 2.08 | 940.43 | 13.96 | 0.61 | a | b | b |
| Asparagine | 80.3 | 0.73 | 0.27 | 51.12 | 0.102 | 0.046 | a | b | b |
| Proline | 2.31 | 0.11 | 0.08 | 1.50 | 0.090 | 0.017 | a | b | b |
| Arginine | 2.8 | 0.53 | 0.21 | 2.81 | 0.26 | 0.056 | a | a | a |
| b-aminoisobutyrate | 0.36 | 0.01 | 0.01 | 0.37 | 0.0004 | 0 | a | b | b |

| | | | | | | | | | | |
|---|---|---|---|---|---|---|---|---|---|---|
| | b-Alanine | 7.63 | 0.11 | 0.07 | 6.10 | 0.036 | 0.017 | a | b | b |
| **Organic acids** | **Gluconate** | 68.19 | 0.95 | 0.01 | 15.58 | 0.71 | 0.0014 | a | b | c |
| | **Galactonate** | 2.47 | 0.19 | 0.03 | 0.64 | 0.17 | 0.021 | a | b | c |
| | **Fumarate** | 2.42 | 0.23 | 0.05 | 1.33 | 0.15 | 0.014 | a | b | c |
| | **Shikimate** | 0.43 | 0.06 | 0.01 | 0.48 | 0.032 | 0.0042 | a | b | c |
| | **Glycerate** | 1.73 | 0.26 | 0.09 | 1.51 | 0.25 | 0.033 | a | b | c |
| | Citrate | 6.81 | 0.06 | 0.01 | 3.91 | 0.014 | 0.0004 | a | b | b |
| | Malate | 16.07 | 0.22 | 0.05 | 4.13 | 0.097 | 0.018 | a | b | b |
| | Succinate | 6.06 | 0.15 | 0.06 | 3.42 | 0.0175 | 0.0053 | a | b | b |
| | Quinate | 0.01 | 0.01 | 0 | 0.0087 | 0.0004 | 0.0017 | a | ab | b |
| | Glycolate | 0.86 | 0.1 | 0.09 | 1.06 | 0.013 | 0.029 | a | b | b |
| | Citramalate | 0.17 | 0.07 | 0 | 0 | 0.017 | 0 | a | a | a |

| | | | | | | | | | | |
|---|---|---|---|---|---|---|---|---|---|---|
| | Salicylate | 0.07 | 0.42 | 0.19 | 0.071 | 0.13 | 0.13 | a | a | a |
| | 2-Oxoglutarate | 0.88 | 0.06 | 0.01 | 0.88 | 0.006 | 0.0014 | a | b | b |
| **Sugars** | **Fructose** | 71.4 | 7.24 | 1.17 | 66.39 | 7.41 | 0.22 | a | b | c |
| | **Galactose** | 11.84 | 1.41 | 0.44 | 8.03 | 0.803 | 0.15 | a | b | c |
| | **Glucose** | 169.59 | 1.98 | 0.59 | 129.67 | 1.35 | 0.22 | a | b | c |
| | **Maltose** | 0.7 | 0.55 | 0.08 | 0.37 | 0.502 | 0.037 | a | a | b |
| | Rhamnose | 0.3 | 0.19 | 0.09 | 0.33 | 0.104 | 0.049 | a | ab | b |
| | Mannose | 1.05 | 2.4 | 0.17 | 0.71 | 0.170 | 0.10 | a | b | b |
| | Xylose | 0.82 | 0.2 | 0.21 | 0.63 | 0.169 | 0.11 | a | b | b |
| | Sucrose | 1.15 | 0.3 | 0.25 | 0.96 | 0.102 | 0.085 | a | a | a |
| | Trehalose | 0.13 | 0.07 | 0.07 | 0.11 | 0.060 | 0.017 | a | a | a |
| **Phosphate** | Phosphate | 209.6 | 5.2 | 0.8 | 182.60 | 0.73 | 0.79 | a | b | b |

| | | | | | | | | | | |
|---|---|---|---|---|---|---|---|---|---|---|
| | Glucose-6-P | 0.78 | 0.02 | 0 | 0.34 | 0.0004 | 0.0004 | a | b | b |
| | Fructose-6-P | 0.27 | 0 | 0 | 0.22 | 0 | 0 | a | b | b |
| **Others** | Erythritol | 0.25 | 0.65 | 0.52 | 0.18 | 0.61 | 0.34 | a | ab | b |
| | Putrescine | 0.95 | 0.2 | 0.28 | 0.68 | 0.20 | 0.065 | a | b | b |
| | Ethanolamine | 4.18 | 0.19 | 0.15 | 4.63 | 0.14 | 0.054 | a | b | b |
| | Arabitol | 0.12 | 0.15 | 0.07 | 0.091 | 0.051 | 0.043 | a | a | a |
| | Glycerol | 70.58 | 137.53 | 36.08 | 55.76 | 29.70 | 30.28 | a | a | a |
| | **Myo-Inositol** | 11.51 | 2.8 | 0.67 | 8.01 | 3.09 | 0.303 | a | b | c |

[a] n= 12
[b] Different letters indicate significantly different content with p<0.05 using a non-parametric Kruskal-Wallis statistical test.
[c] Metabolite contents presented in Figure 4B that are significantly lower in the guttation fluid compared to the xylem sap are indicated with names in bold.

**Table S5:** Name and sequence of oligonucleotides used in this study.

| **Name** | **Gateway primers (5'-3')**[a] |
|---|---|
| AT3G16670_prom_F | <u>GGGGACAAGTTTGTACAAAAAAGCAGGCT</u>GTGAATCTGTTTCAAAATCTATGAG |
| AT3G16670_prom_R | <u>GGGGACCACTTTGTACAAGAAAGCTGG</u>GTCTTTTTTGGATTACTTGTATATGAAAC |
| AT3G05730_prom_F | <u>GGGGACAAGTTTGTACAAAAAAGCAGGCT</u>TCACGATTGTTTCCTCCCTATAC |
| AT3G05730 prom_R | <u>GGGGACCACTTTGTACAAGAAAGCTGG</u>GTCTTTGGAAGTTTTTGCTTTCTTTTC |
| AT1G62510_Prom_F | <u>GGGGACAAGTTTGTACAAAAAAGCAGGCT</u>TCGGGTTGATAGATACTGGTACG |
| AT1G62510_Prom_R | <u>GGGGACCACTTTGTACAAGAAAGCTGG</u>GTCTGTTAATCTCACTTTTGGTTATAGAGG |
| AT1G56710_prom_F | <u>GGGGACAAGTTTGTACAAAAAAGCAGGCT</u>TCGTTACTTCGGCTATTTTGCTAATC |
| AT1G56710_prom_R | <u>GGGGACCACTTTGTACAAGAAAGCTGG</u>GTCTTTTTGTGAATGTCTTAGGAG |

[a] *attB* sites underlined
